# Modeling microbial metabolic trade-offs in a chemostat

**DOI:** 10.1101/664698

**Authors:** Zhiyuan Li, Bo Liu, Sophia Hsin-Jung Li, Christopher G. King, Zemer Gitai, Ned S. Wingreen

## Abstract

Microbes face intense competition in the natural world, and so need to wisely allocate their resources to multiple functions, in particular to metabolism. Understanding competition among metabolic strategies that are subject to trade-offs is therefore crucial for deeper insight into the competition, cooperation, and community assembly of microorganisms. In this work, we evaluate competing metabolic strategies within an ecological context by considering not only how the environment influences cell growth, but also how microbes shape their chemical environment. Utilizing chemostat-based resource-competition models, we exhibit a set of intuitive and general procedures for assessing metabolic strategies. Using this framework, we are able to relate and unify multiple metabolic models, and to demonstrate how the fitness landscape of strategies becomes intrinsically dynamic due to species-environment feedback. Such dynamic fitness landscapes produce rich behaviors, and prove to be crucial for ecological and evolutionary stable coexistence in all the models we examined.

## INTRODUCTION

The way microbes respond to and shape their local environment influences their community structure (Callahan, Fukami, & Fisher, 2014). Such microbe-environment interactions depend on the allocation strategies of cells, i.e., how a cell allocates its internal resources into various cellular functions, such as transport, assimilation, reproduction, motility, maintenance, etc. (Bachmann, Bruggeman, Molenaar, dos Santos, & Teusink, 2016). Within a microbial cell, energy and biomass are limited, and trade-offs always exist in allocating these valuable internal resources into the various functions required for cell growth. Therefore, the growth rate of cells cannot increase without bound. Rather, evolution acts on cells’ internal resource allocation to optimize growth and survival (S. Goyal, Yuan, Chen, Rabinowitz, & Wingreen, 2010; Liebermeister et al., 2014). To this end, in response to environmental changes, microbes rapidly adjust their metabolic strategies. For example, the yeast *Saccharomyces cerevisiae* switches from fermentation to respiration upon glucose depletion (Zaman, Lippman, Zhao, & Broach, 2008), and *Escherichia coli* exhibits drastic differences in ribosome content between different nutrient conditions (Li et al., 2018; Scott, Gunderson, Mateescu, Zhang, & Hwa, 2010). Moreover, in laboratory long-term evolution studies of microbes, adapative mutations consistently emerge that reshape metabolism (Bachmann, Molenaar, dos Santos, & Teusink, 2017; Bajic & Sanchez, 2019; Blount, Barrick, Davidson, & Lenski, 2012; Long & Antoniewicz, 2018). Such short-term and long-term adjustments of metabolic strategies presumably confer fitness benefits, and it is important to map metabolic strategies onto these benefits to better understand the regulation and evolution of microbial metabolism.

The convergence towards steady state makes the chemostat an ideal experimental system to culture microorganisms and investigate their metabolic status (Wides & Milo, 2018; Ziv, Brandt, & Gresham, 2013). In a chemostat, fresh nutrients are supplied at a constant rate, while medium with cells is removed at the same rate to maintain constant volume. The metabolite concentrations in the chemostat constitute the chemical environment directly perceived by cells, and determine their growth rates. Importantly, cells also shape this environment through their consumption and secretion of metabolites. One advantage of a chemostat is the automatic convergence of cellular growth rates towards the controlled dilution rate. This convergence occurs through negative feedback between microbes and their environment: the higher the population, the worse the chemical environment, and the slower the growth rate. As a result (provided the nutrient supply allows for faster-than-dilution growth to prevent “washout”), the cells in the chemostat will reach the steady-state population that sustains growth at the dilution rate (De Leenheer, Levin, Sontag, & Klausmeier, 2006; Smith & Waltman, 1995). This stabilization of the cellular growth rate at the controlled dilution rate facilitates precise characterization of cellular physiology in a constant environment. However, it also imposes challenges in quantitatively understanding the advantages and disadvantages of various metabolic strategies: if all metabolic strategies lead to identical growth rates in a chemostat, how should we evaluate whether one strategy is “better” or “worse” than another? If strategies can be compared, are there “best” strategies, and how do these optima shift as the experimental conditions such as nutrient-supply concentrations and dilution rates change?

If we evaluate metabolic strategies by the outcome of competition between species, many insights can be gained from theoretical ecology. For example, resource-competition models have provided a simple context to explore competition dynamics in chemostat-like ecosystems such as lakes and rivers (Smith & Waltman, 1995). In such models, species interact only indirectly via consumption (and sometimes production) of a common pool of nutrients. A steady state can be reached if the species present can shape the nutrient concentration to support a growth rate equal to their dilution or death rate (Tilman, 1982). Resource-competition models underpin many ecosystem theories including contemporary niche theory as pioneered by MacArthur (MacArthur, 1970), popularized by Tilman (Tilman, 1980, 1982), and extended by Chase and Leibold (Chase & Leibold, 2003). A central component of contemporary niche theory is a graphical approach, generally consisting of three components: zero net growth isoclines (ZNGIs) in chemical space, an impact vector representing a species’ nutrient consumption, and a supply point to describe the external resource supply (Koffel, Daufresne, Massol, & Klausmeier, 2016). This graphical approach is a powerful and intuitive way of assessing the outcome of competition, yet it is not yet commonly utilized in understanding microbial metabolic strategies with trade-offs.

Resource-competition models focusing on various aspects of cellular metabolism vary in their assumptions regarding species-environment interactions, and can lead to diverse results for community structure and population dynamics. In a model where species compete for essential resources, different nutrient requirements can produce intrinsically oscillatory or even chaotic dynamics (Huisman & Weissing, 1999, 2001). Alternatively, cross-feeding (Goldford et al., 2018; Pfeiffer & Bonhoeffer, 2004) can promote stable coexistence, while preferential nutrient utilization (A. Goyal, Dubinkina, & Maslov, 2018) can lead to multistability. With metabolic trade-offs, a model in which growth rate is additive in imported nutrients self-organizes to a state of unlimited stable coexistence (Posfai, Taillefumier, & Wingreen, 2017), while another model with convertible essential nutirents also allows evolutionarily stable coexistence but with a limited number of species (Taillefumier, Posfai, Meir, & Wingreen, 2017). This large variety of models and the richness of possible behaviors raises the question of unification: is there a simple framework that consolidates this diverse group of models into one easily understandable picture?

Continuity of the strategy space adds another layer of complexity in characterizing the “best” metabolic strategy or strategies. With infinite possibilities for allocating cellular resources, how should one pinpoint the optimal ones? Adaptive dynamics in ecological theory, also known as evolutionary invasion analysis, provides valuable guidance (Metz, 2012). This mathematical framework adtresses the long-term evolution of traits in asexually reproducing populations by quantifying the fitness of each “trait” as a function of population composition. In this framework, “invasion fitness” is defined as the net-growth rate of a new variant when it is introduced into the resident population in an infinitesimally small amount, and a population allowing only non-positive invasion fitness for any new variant is considered to be “evolutionarily stable” (Doebeli, 2002; Ispolatov, Madhok, & Doebeli, 2016; Rueffler, Van Dooren, & Metz, 2004). Such an evolutionarily stable point is valuable for defining optimal strategies, as a community adopting the most suitable metabolic strategies should not be invasible by any other strategies. As microbes in nature frequently experience environmental fluctuations, it is important to understand whether and how such “optimal metabolic strategies” change with external conditions. Nevertheless, in the standard modeling framework of adaptive dynamics, the environment is implicit, and species directly act on each other without the realistic constraints imposed by competition for resources. Combining the concept of invasion fitness with explicit competition for resources, under the assumption of metabolic trade-offs, could therefore bring new insights into microbial metabolic strategies and community assemblies.

In this work, we present a mathematical framework for analyzing competition for resources among various metabolic strategies in a chemostat setting. We combine and extend the graphical tools from resource-competition theory and the invasion-fitness approach from adaptive dynamics to relate and unify multiple models for microbial metabolic trade-offs. This combination provides an intuitive scheme to evaluate strategies under various external conditions. The center of the framework is the role of species in creating their own environment. Firstly, the chemical environment shaped by an endogenous species through growth and consumption can be inviting or prohibiting to an invader species, depending on the geometric relationship between the “zero net growth surface” of the invader and the environment created by the endogenous species. This geometry-dependent fitness leads to a general “rule of invasion” to compare pairs of strategies. In evaluating a continuum of strategies, the ensemble of their invasion fitnesses form a landscape, whose shape depends on the chemical environment. This intrinsically dynamic fitness landscape allows for the intransitivity of fitness (Soliveres et al., 2015). We demonstrate how such intransitivity can lead to rich ecosystem dynamics, including mutual invasion, multistability, and oscillations, and how all of these behaviors can be simply related via a graphical representation and the dynamic fitness landscape. Moreover, from the environment-dependent fitness landscape, we can define non-invasible/optimal metabolic strategies – namely, one or more strategies that construct a fitness landscape that places themselves on the top. The mathematical framework we present confers several advantages. Firstly, it separates the effect of changing dilution rate and of varying supply concentrations, facilitating quantitative interpretation as well as predictions for chemostat experiments. Secondly, it establishes an intuitive mapping from various metabolic models to population dynamics. Additionally, it reveals long-term implications, particularly in clarifying the general conditions for coexistence on both ecological and evolutionary time scales.

## Results

### Metabolic trade-offs and metabolic strategies

As discussed above, microorganisms need to allocate their limited internal resources into different cellular functions. In our models, we use *α*_*j*_ to denote the fraction of internal resources allocated to the *j*-th metabolic function, with 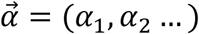 representing a metabolic strategy. An exact metabolic trade-off is assumed, such that ∑_*j*_ *α*_*j*_ = 1. All possible values of 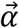 define a continuous spectrum of metabolic strategies, which we name the strategy space. One major goal of our work is to construct a general and intuitive framework for evaluating strategies for a broad range of different metabolic models and experimental conditions.

### Graphical representation of species creating their own chemical environment

One way to evaluate metabolic strategies is by comparing the competitiveness of “species” with fixed strategies in chemostat-type resource-competition models. In an idealized model of a chemostat (Fig 1A), *p* types of nutrients are supplied at rate *d* and concentrations 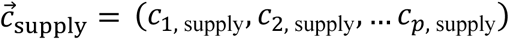, meanwhile cells and medium are diluted at the same rate *d*. The chosen values of *d* and 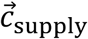 constitute the “external condition” for a chemostat. Nevertheless, the chemical environment that directly impacts cells is the metabolite concentration inside the chemostat, 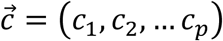, which influences the intracellular metabolite concentrations 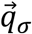 and the growth rate *g*_*σ*_ of the species *σ*. Accordingly, the biomass density *m*_*σ*_ of species *σ* adopting strategy 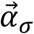 in the chemostat obeys:

**Figure 1.**
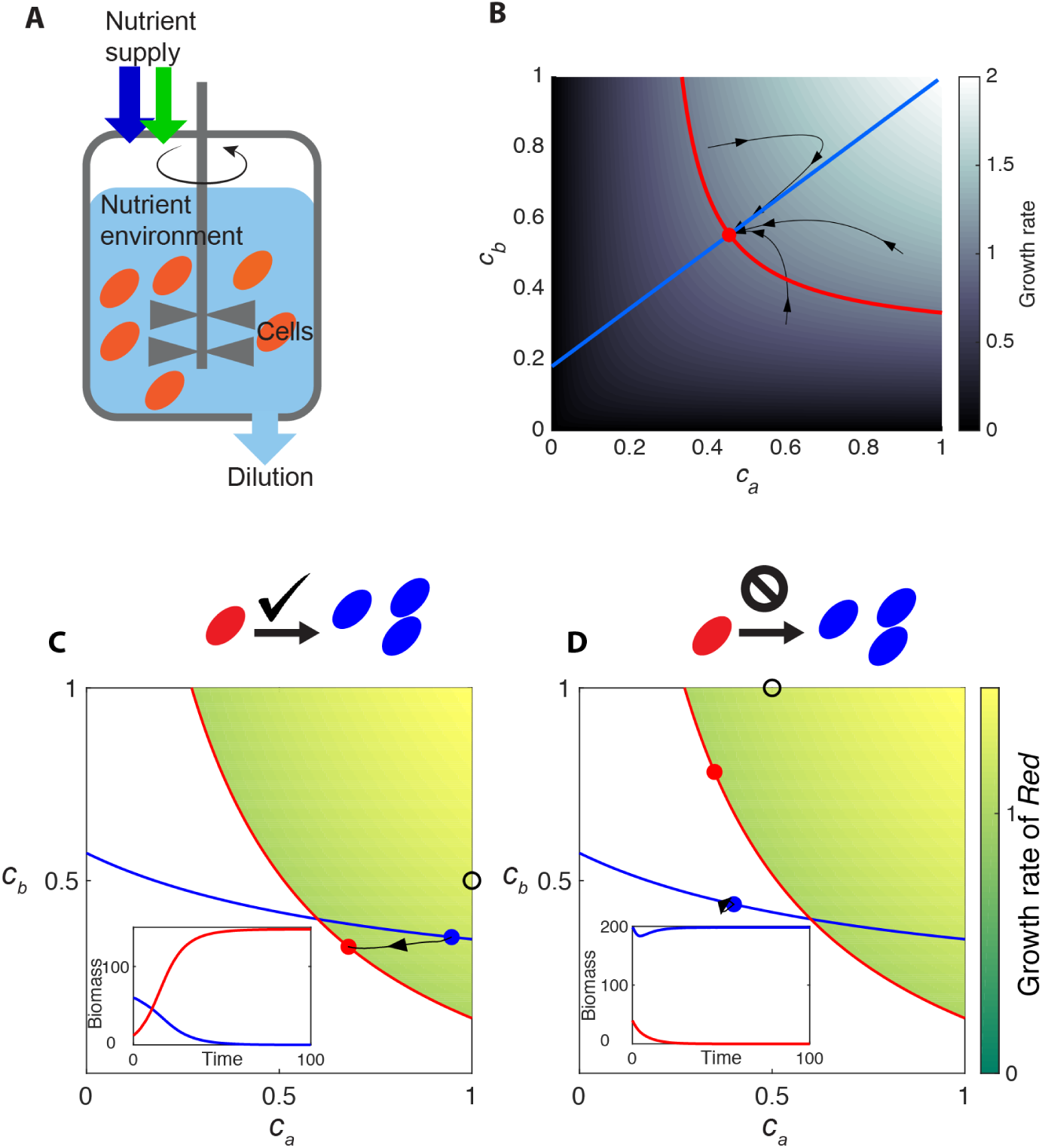
Chemostat behavior represented in chemical space. A. Schematic diagram of a chemostat occupied by a single microbial species. In the well-mixed medium (pale blue) of a chemostat, cells (orange ellipses) consume nutrients and grow. An influx of nutrients with fixed concentrations (blue and green arrows) is supplied at the same rate as dilution, keeping the medium volume constant. B. Visual representation of how a species creates its own chemostat environment. Background color indicates the growth rate of cells as a function of metabolite concentrations *c*_*a*_ and *c*_*b*_, with the growth contour shown by the red curve. The flux-balance curve is shown in blue. Black curves with arrows show the time trajectories of chemostat simulation. C. Example of successful invasion of species *Blue* by species *Red*. A small amount of species *Red* is introduced to a steady-state chemostat of species *Blue*. Growth contours and steady-state environments of species *Blue* and species *Red* are shown as curves and dots in the corresponding colors (colored background indicates the “invasion zone” of *Red*, and represents the growth rate of *Red* in this zone). The supply condition is marked by a black circle. Black curves with arrows show the time trajectory of the invasion in chemical space. Inset: time course of species biomass in the chemostat during the invasion. D. Same as (C), except that because the supply condition (black circle) is different, the attempted invasion by species *Red* is unsuccessful.

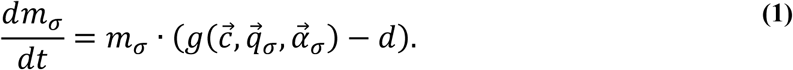

The concentration *c*_*i*_ of the *i*-th nutrient is a variable, influenced by its rate of consumption *I*_*i*_ per cell volume. In a chemostat occupied by a single species *σ*, the changing rate of *c*_*i*_ satisfies

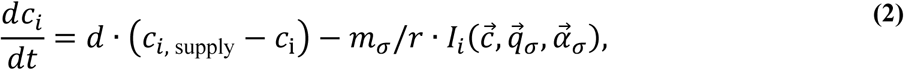

where *r* is a constant representing the biomass per cell volume. (If the volume of the chemostat is *V*_chemostat_ and the total volume of cells is *V*_cells_, the import flux of the *i*-th nutrient *V*_cells_ · *I*_*i*_ implies a rate of change of concentration inside cells of *I*_*i*_ and a corresponding rate of change of the concentration in the chemostat of *V*_cells_ /*V*_chemostat_ · *I*_*i*_ = (*m* · *V*_chemostat_/*r)*/*V*_chemostat_ · *I*_*i*_ = *m*/*r* · *I*_*i*_.). A negative value of *I*_*i*_ corresponds to secretion of the metabolite from cells into the environment.

In this manuscript, we define 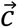 as the “chemical environment”, and all possible values of 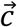 constitute the “chemical space”. Within a cell, the concentration of metabolites is influenced by uptake/secretion rates, and influences the growth rate. Different metabolic models assume different forms for such influences, and we use 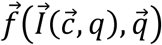 to represent the rate of change in 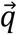:

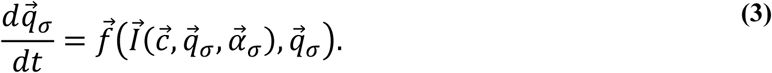

Eqs. (1)-(3) represent a general chemostat model with a single species. The simplicity of the chemostat has inspired many theoretical studies of resource competition. Different model assumptions about how species grow and consume nutrients have produced a variety of intriguing behaviors and conclusions. However, the origins of these differences are not always simple to discern. To facilitate the evaluation of metabolic strategies, we next present a graphical representation that allows ready visualization of the feedback between species and the environment in a chemostat.

The steady-state environment created by species *σ* can be obtained by setting Eqs. (1)-(3) to zero. First, when Eq. (3) is equal to zero, 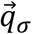 can be solved as a function of 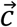, reducing 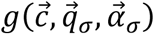 and 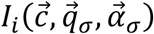 to functions fully dependent on the variable 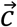, namely 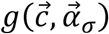 and 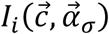. Graphically, the steady state created by a single species can be visualized by the intersection of two nullclines, derived from Eq. (1) and Eq. (2), respectively (details in Methods).

Next, setting Eq. (1) to zero leads to a *p*-dimensional version of the ZNGI, which we name the “*growth contour”*. For a given metabolic strategy 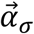, the growth-rate function 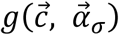 maps different points in the chemical space onto varying growth rates (background color in Fig 1B). At steady state, the relation *dm*_*σ*_/*dt* = 0 (Eq. (1)) requires the growth rate to be exactly equal to the dilution rate (assuming nonzero cell density). Therefore, the contour in chemical space satisfying 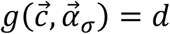 indicates all possible environments that could support a steady state of the strategy 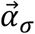 (red curve in Fig 1B). This contour reflects how the chemical environment determines cell growth.

Thirdly, the nullcline derived from Eq. (2) reflects the impact of cellular metabolism on the chemical environment. At steady state, the nutrient influx should be equal to the summation of dilution and cellular consumption. When Eq. (2) is set to zero, varying values of cell density *m* lead to different 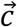 (Eq. (S5)), constituting a one-dimensional “flux-balance curve” in chemical space (purple, cyan, and blue curves in Fig 1B).

Finally, the steady-state chemical environment 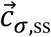 created by the species *σ* is the intersection of the growth contour and the flux-balance curve (Fig 1B, red dot). Changes in the chosen conditions *d* and 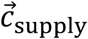 influence the shapes of the growth contour and the flux-balance curve separately, enabling clear interpretation of chemostat experiments under varied conditions (details in Methods). Sometimes for a given steady-state environment 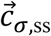, one would like to derive the supply concentrations that ultimately lead to this environment. Setting Eq. (2) to zero and fixing 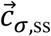, varying values of cell density *m* lead to a straight line in the space of supply concentrations, which we call the “supply line” (see Methods for details). Despite the fact that the supply space and the chemical space are distinct, they share the same units of concentration in each dimension. Therefore, for ease of visualization we typically show supply lines along with other features in the same nutrient space (Eq. (S6), black dashed line in Fig S1).

### Rule of invasion, and the environment-dependent fitness landscape

We use the outcome of competition between species to evaluate metabolic strategies, assuming each species *σ* adopts a fixed strategy 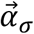. We first focus on the outcome of invasion, which corresponds to the “invasion fitness” in adaptive dynamics: here, “invasion” is defined as the introduction of a very small number of an “invader” species to a steady-state chemostat already occupied by a set of “indigenous” species, and the “invasion growth rate” is quantified by the growth rate of the invader species at the moment of introduction. Unlike adaptive dynamics, the invasion growth rate is evaluated with respect to the chemical environment created by the indigenous species, rather than with respect to the population composition of the indigenous species.

In the chemical space, the outcome of an invasion can be summed up by a simple geometric rule, as demonstrated in Fig 1C and D. The growth contour of the invader (species *Red*) separates the chemical space into two regions: an “invasion zone” where the invader grows faster than dilution (green-colored region in Fig 1C and D), and “no-invasion zone” where the invader has a growth rate lower than dilution. If the steady-state environment constructed by the indigenous species (species *Blue*) is located within the invasion zone of the invader, the invader will initially grow faster than dilution. Therefore, the invader will expand its population and the invasion will be successful (Fig 1C). By contrast, if the steady-state chemical environment created by the indigenous species lies outside of the invasion zone, the invasion will be unsuccessful (Fig 1D, same species as in Fig 1C but with a different supply condition, and therefore a different steady state). (See Methods for details.)

In the steady-state environment created by the indigenous species, each strategy 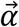 has an invasion growth rate. We define an environment-dependent “fitness landscape” as the relation between the invasion growth rate and the metabolic strategy of invaders (Eq. (S8)-(S9), see Methods for details). Different indigenous species can create different chemical environments, and thus give rise to different shapes of the fitness landscapes.

### Mutual invasion, a flat fitness landscape, and unlimited coexistence

The rule of invasion allows for easy assessment of competition dynamics. The emergence of complex dynamics generally requires that competitiveness be non-transitive. For example, if species *Red* can invade species *Blue*, that does not mean *Blue* cannot invade *Red*. Such mutual invasibility can be observed in substitutable-resource metabolic models, with a simple version illustrated in Fig 2A: two substitutable nutrients *a* and *b*, such as glucose and galactose, contribute linearly to biomass increase. Since a substantial investment of protein and energy is required for nutrient uptake, the model assumes a trade-off between the allocation of internal resources to import either nutrient. Specifically, a fraction of resources *α*_*a*_ is allocated to import *a* and a fraction (1 − *α*_*a*_) to import *b.* As shown in Fig 2B, while the steady-state environment created by *Blue* is located within the invasion zone of *Red*, the steady-state environment created by *Red* is also located within the invasion zone of *Blue*. According to the rule of invasion, each species can therefore invade the steady-state environment created by the other. In the face of such successful invasions, the only possible stable chemical environment for this system is at the intersection of two growth contours, where the two species can coexist.

**Figure 2.**
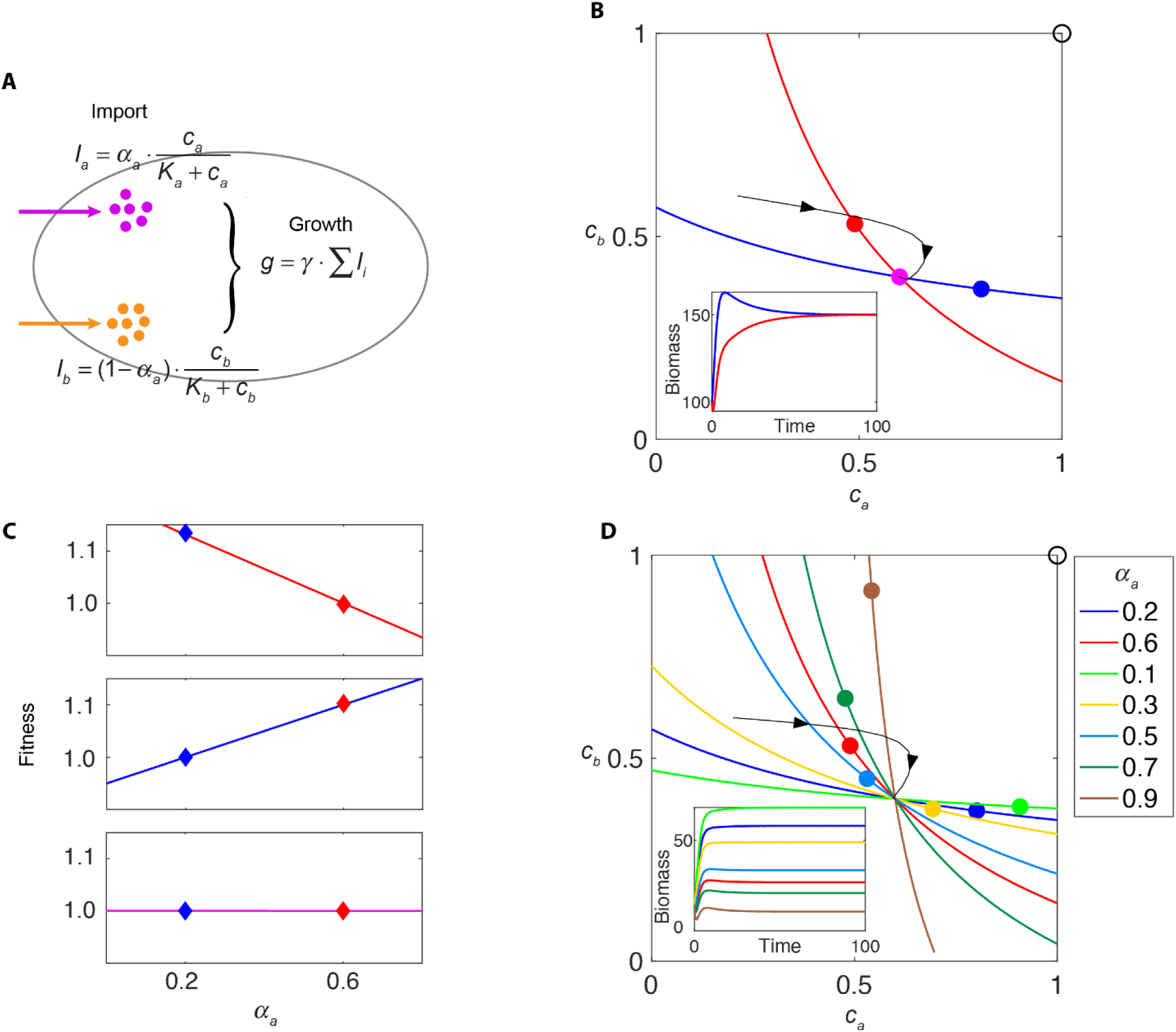
Metabolic models with substitutable nutrients and coexistence in a chemostat. A. Example of a metabolic model with a trade-off in allocation of internal resources for import of two substitutable nutrients, with both nutrients contributing additively to growth. Species *Red* and species *Blue* allocate resources differently (indicated by parameter *α*_*a*_, see Methods). B. Growth contours and the steady-state environments created by *Red* or *Blue* alone, under the supply condition shown by the black circle. Black curve with arrows shows a trajectory in chemical space. Purple dot indicates the steady-state environment created by *Red* and *Blue* together. Lower inset: time course of species biomass. C. From top panel to bottom panel: the fitness landscape created by *Red* alone (for the red dot in (B)), created by *Blue* alone (for the blue dot in (B)), and created by both species (for the purple dot in (B)). Diamonds mark the locations of *Red* and *Blue* strategies and their corresponding fitness in each fitness landscape. D. Growth contours and the species-specific steady-state environments for seven different species alone, under the supply condition shown by the black circle. Black curve with arrows shows a trajectory in chemical space. Lower inset: time course of species biomass in the chemostat.

The environment-dependent fitness landscape readily explains this coexistence: In the steady-state environment created by *Red* (*α*_*a*_ = 0.6), strategies with smaller *α*_*a*_ have higher fitness (Fig 2C, upper panel). In the steady-state environment created by *Blue* (*α*_*a*_ = 0.2), the fitness landscape changes in shape, and strategies with larger *α*_*a*_ have higher fitness (Fig 2C, middle panel). Therefore, each species creates an environment that is more suitable for its competitor, which leads to coexistence.

Typically in resource-competition models, the number of coexisting species cannot exceed the number of resources (Armstrong & McGehee, 1980; Hardin, 1960; Levin, 1970; McGehee & Armstrong, 1977). This conclusion can be understood intuitively from a graphical approach: The steady-state growth of a species can only be achieved on the growth contour of this species. The stable-coexistence of *N* species can thus only occur at the intersection of their *N* corresponding growth contours. Generally, an *N*-dimensional chemical space allows a unique intersection of no more than *N* surfaces, and the diversity of species is therefore bounded by the number of metabolites in the environment. This theoretical restriction on biodiversity, made formal as the “competitive exclusion principle”, contradicts the tremendous biodiversity manifested in the real world (Friedman, Higgins, & Gore, 2017; Goldford et al., 2018; Maharjan, Seeto, Notley-McRobb, & Ferenci, 2006). There have been a multitude of theoretical efforts to reconcile this contradiction (Huisman & Weissing, 1999, 2001)(Goldford et al., 2018; Pfeiffer & Bonhoeffer, 2004)(Beardmore, Gudelj, Lipson, & Hurst, 2011; Taillefumier et al., 2017).

The mutual invasibility in linear-summation metabolic models with trade-offs presents one possibility resolution of the competitive-exclusion paradox. For the environment co-created by *Blue* and *Red* (Fig 2B, purple dot), the fitness landscape becomes flat (Fig 2C, bottom panel): species with any metabolic strategy will grow at the same rate as dilution in this environment. Therefore, in this system, once a pair of species with a mutual-invasion relationship construct the chemical environment together, all species become effectively neutral and can coexist indefinitely (see Methods for details). This graphical approach to mutual invasion and the flat fitness landscape provide an intuitive representation of species self-organizing to a state of unlimited coexistence beyond competitive exclusion.

### Rock-paper-scissor invasion loop and oscillation

Resource-competition models focusing on various aspects of cellular metabolism vary in their assumptions regarding 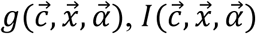, and 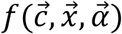, and can lead to diverse results for community structure and coexistence. However, the above general “rule of invasion” allows us to treat these divergent resource-competition models in a unified framework. In the following example, we utilized a metabolic model slightly different from that of Fig 2, to show that a dynamic fitness landscape is indispensable for coexistence. In the metabolic model shown in Fig 3A, three substitutable nutrients, *a, b*, and *c*, contribute additively to cell growth. In this three-dimensional chemical space, the growth contour for each species is a two-dimensional surface (Fig 3B). In addition to requiring enzymes to import the raw forms of these nutrients as in the model of Fig 2A, enzymes are also required to convert the imported raw materials into biomass. In this model, a six-element 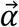 is required to describe the metabolic strategy, and there is the possibility of “mismatches” in the fraction of internal resources allocated to import and to convert the same nutrient. Such mismatches can produce a “rock-paper-scissor” invasion loop (Fig S3A): In the environment created by species 1, species 2 has a higher fitness but not species 3; therefore species 2 can invade species 1 and establish its own environment; however, this environment lies within the invasion zone of species 3 (Fig 3B) but not of species 1, therefore species 3 subsequently invades; then in turn, species 3 create an environment where species 1 has the highest fitness. Such a loop of invasions leads to oscillatory population dynamics (Fig 3C, upper panel), with an ever-changing fitness landscape (Fig 3C, lower panel).

**Figure 3.**
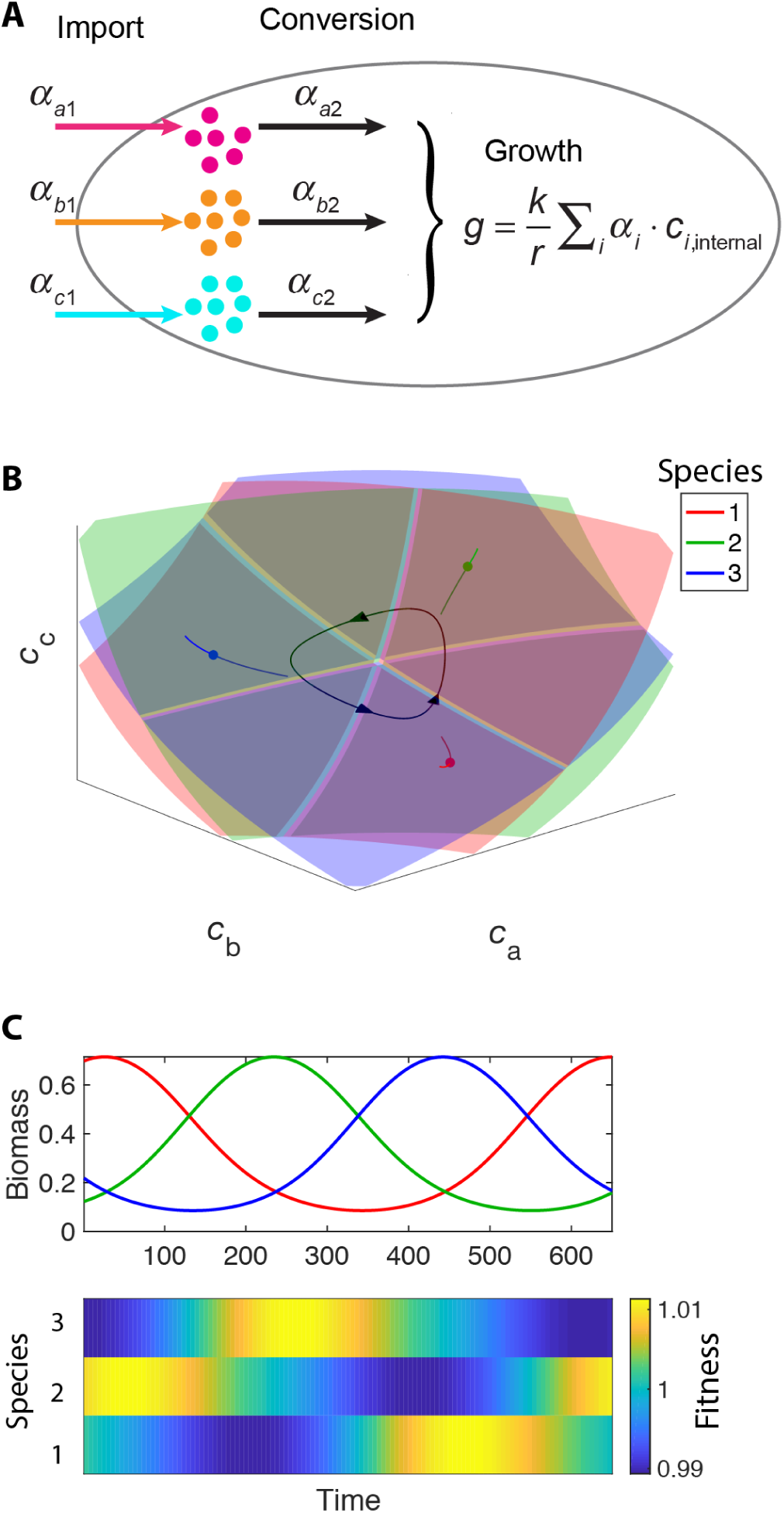
Rock-paper-scissors oscillations. A. Example of a metabolic model with a trade-off in allocation of internal resources for import and assimilation of three substitutable nutrients, with all three nutrients contributing additively to growth. Species *Red*, species *Blue*, and species *Green* allocate resources differently (see Methods). B. Growth contours (surfaces), flux-balance curves (lines), and steady-state nutrient concentrations (dots) for the three species in a three-dimensional chemical space. Black curves with arrows show the system’s limit-cycle trajectory. C. The top panel shows the time course of species biomass in the chemostat for the limit cycle in (B). The bottom panel shows how the fitness landscape changes with time over one period of the oscillation.

Oscillation and even chaos in resource-competition model have been demonstrated by Huisman et al. (Huisman, van Oostveen, & Weissing, 1999), and shown to allow dynamical coexistence beyond competitive exclusion. The simple model presented here illustrates how oscillation can be understood as a loop of invasion creating an ever-changing fitness landscape.

### Multistability and chain of invasion

When species create environments that are more favorable for their competitors, mutual-invasion and oscillations can occur. Can species create environments that are hostile to their competitors, and if so what will be the consequences?

Figure 4A shows a simple metabolic model with two essential nutrients *a* and *b*, such as nitrogen and phosphorus (see Methods for details). Similar to the model in Fig 2A, the model assumes a trade-off between the allocation of internal resources to import nutrients, so that a resource allocation strategy is fully characterized by the fraction of resources *α*_*a*_ allocated to import nutrient *a*. The growth rate is taken to be the minimum of the two input rates (Odum & Barrett, 1971). As shown in Fig 4B, two species, *Red* and *Blue*, each creates a chemical environment outside of the invasion zone of each other. According to the rule of invasion, neither can be invaded by the other. Therefore, the steady state of the community depends on initial conditions – whichever species occupies the chemostat first will dominate indefinitely. It is worth noting that despite the fact that coexistence is excluded in a single chemostat under this metabolic model, the ability of species to create a self-favoring environment allows the spontaneous emergence of spatial heterogeneity and coexistence in an extended system with multiple linked chemostats (Fig S6). In ecology, the spontaneously emergence of spatial heterogeneity has been shown for species with the capacity to construct their own niches (Levin, 1970; Schertzer, Staver, & Levin, 2015), and the chain of chemostats provides a simple model for such spatial coexistence.

**Figure 4.**
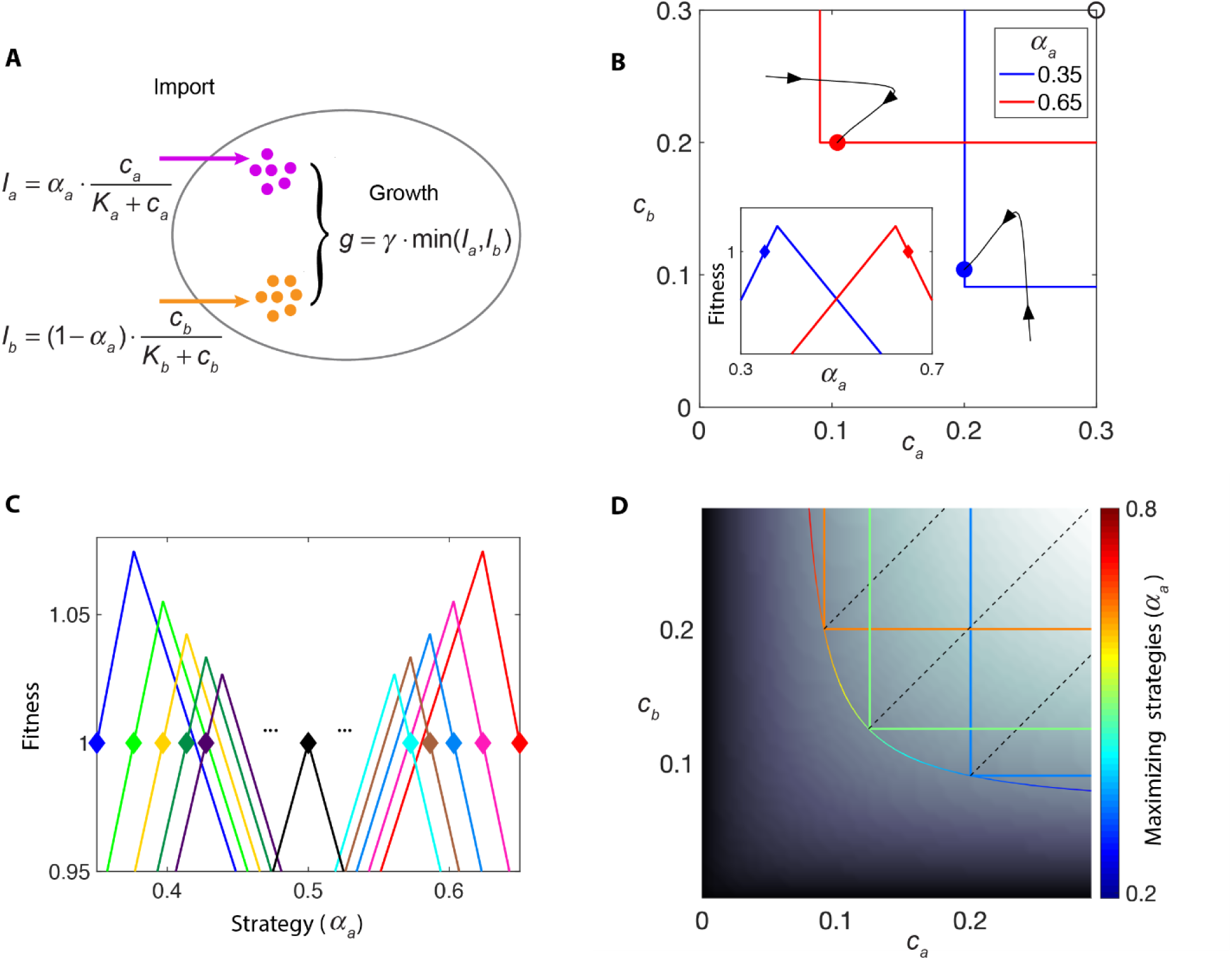
Multistability, chain of invasion, and non-invasible strategy. A. Example of bistability for a metabolic model with a trade-off in allocation of resources for import of two essential nutrients, with the lower of the two import rates determining growth rate. Species *Red* and species *Blue* allocate resources differently (indicated by parameter *α*_*a*_, see Methods). B. Bistability of the system in (A) shown in chemical space. Black curves with arrows show the trajectories of simulations with different initial conditions. Inset: the fitness landscape created by species *Red* or *Blue* alone, with colors corresponding to the steady-state environments shown by colored dots in the main panel. C. A chain of invasion. Fitness landscape created by species with different internal resource allocation strategies (marked by diamond shapes). Starting from species *Blue*, the species having the highest growth rate in the fitness landscape created by the “former” species is chosen. This creates a chain of invasion from *Blue* to *Light Green, Yellow, Deep Green, Deep Purple*, all the way (intermediate processes omitted) to the species *Black*, which places itself on the peak of its own fitness landscape. The same procedure is also performed starting with species *Red*. D. Depiction of non-invasible strategies under different supply conditions. Black-white background indicates the maximal growth rate of model in (A) under each environment, and the contour of maximal growth rates contains different strategies (represented by red-to-blue color). Growth contours of three species adopting one of the “maximizing strategies” are colored by their strategies. The supply conditions allowing these strategies to be “non-invasible” are marked by dashed black lines.

From the perspective of the strategy-growth relationship (Fig 4B, inset), species *Red* (*α*_*a*_ = 0.65) creates a fitness landscape where small *α*_*a*_ is disfavored. Symmetrically, species *Blue* (*α*_*a*_ = 0.35) creates a fitness landscape where large *α*_*a*_ is disfavored. However, neither *Red* nor *Blue* sits on the top of the fitness landscape each one creates (Fig 4C). In the fitness landscape created by *Blue*, a slightly larger *α*_*a*_ (green diamond in Fig 4C) has the highest growth rate. Consequently, species adopting the *Green* strategy can invade *Blue*. Nevertheless, species *Green* is not on the top of its own fitness landscape as an even larger *α*_*a*_ (yellow diamond in Fig 4C) maximizes the growth rate in the environment created by *Green*. A series of replacements by the fastest-growing species in the environment created by the former species creates a chain of invasion.

In this particular model after four steps of replacement, bistability appears. The species with *α*_*a*_ marked by *Deep Purple*, which is reached by the chain of invasion going from *Blue*, to *Green*, to *Yellow*, to *Deep Green*, cannot invade the original species *Blue*. A similar relationship holds between *Cyan* and *Red*. This phenomenon highlights the difference between ecological stability and evolutionary stability: Ecologically, as both *Blue* and *Deep Purple* create a fitness landscape where the other species grows slower than dilution, they constitute a bistable system. However, evolutionarily, “mutants” with slightly larger *α*_*a*_ can invade *Blue,* bringing the system towards *Deep Purple* until bistability collapses.

### Non-invasible strategies

In this model, with symmetric parameters, the only evolutionarily stable strategy is *α*_*a*_ = 0.5 (black diamond in Fig 4C). This is the only strategy that locates itself on the top of the fitness landscape it creates, and therefore cannot be invaded by any other species. This simple model demonstrates a general definition of optimal (aka evolutionarily stable or non-invasible) strategies: those strategies that create a fitness landscape which places themselves on the top (Eq. (S10)).

A chemical environment defines a fitness landscape, and the steady-state chemical environment created by species is influenced by supply condition, dilution rate, and the details of cell metabolism. Therefore, different chemostat parameters and different metabolic models lead to different optimal strategies. In the following, we described a generally applicable protocol for obtaining the non-invasible strategies, using the metabolic model in Fig 4A as the example (Fig 4D, details in Methods): First, under a chemical environment 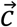, the maximal growth rate 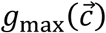 (background color in Fig 4D) and the corresponding resource allocation strategy 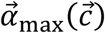 can be obtained analytically or via numerical search through the strategy space (Eq. (S11)). 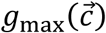 and 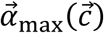 are independent of the chemostat parameters 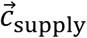 and *d*. Second, the maximal growth contour for dilution rate *d* is defined as all chemical environments 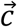 that support a maximal growth rate of *d* (Eq. (S12)). Different maximizing strategies 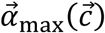 exist at different points of the maximal growth contour, as shown by the colors of the curve in Fig 4D. By definition, the maximal growth contour envelops the growth contour of any single strategy, and chemical environments on the maximal growth contour are outside of the invasion zone of any strategy. Therefore, if a species is able to create a steady-state environment on the maximal growth contour, it cannot be invaded. Finally, different 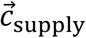 form different maximal flux-balance curves (Eq. (S14)), which intersect with the maximal growth contour at one point 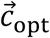. Species 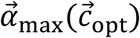 that adopt the maximizing strategy at 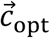 create the environment 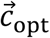, and are therefore immune to invasion. Under different 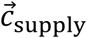, different species become non-invasible (orange, green, and blue growth contours in Fig 4D), and the supply lines emanating from different points on the maximal growth contour indicate the supply conditions for which the corresponding strategies are optimal.

### Evolutionarily stable coexistence

Given *d* and 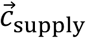, the maximal growth contour and the maximal flux-balance curve are unique, therefore there is only one 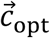. Does the uniqueness of 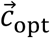 imply a single evolutionarily stable species? Or is coexistence still possible even in the face of evolution? In a recent work (Taillefumier et al., 2017), this question was addressed by modeling a population of microbes competing for steadily supplied resources. Through *in-silico* evolution and network analysis, the authors found that multiple species with distinct metabolic strategies can coexist as evolutionarily-stable co-optimal consortia, which no other species can invade.

Using a simplified version of Taillefumier et al.’s model (Fig 5A), we employ the graphical approach to help identify the requirements for such evolutionarily-stable coexistence and the role of each species in supporting the consortium. In this model, at the cost of producing the necessary enzymes, cells are not only able to import external nutrients, but can also convert any one of the internal nutrients into any other. Meanwhile, nutrients passively diffuse in and out of the cell. The internal concentrations of nutrient *a* and nutrient *b* are both essential for cell growth (see Methods for detail). Therefore, metabolic trade-offs in this system have four elements: the fraction of internal resources allocated to import nutrient *a* (*α*_*a*_) or nutrient *b* (*α*_*b*_) and/or convert one nutrient into another (*α*_*ab*_ converts internal *b* into *a,* and *α*_*ba*_ converts internal *a* into *b*). Each species is defined by its internal resource allocation strategy 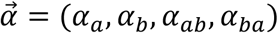.

**Figure 5.**
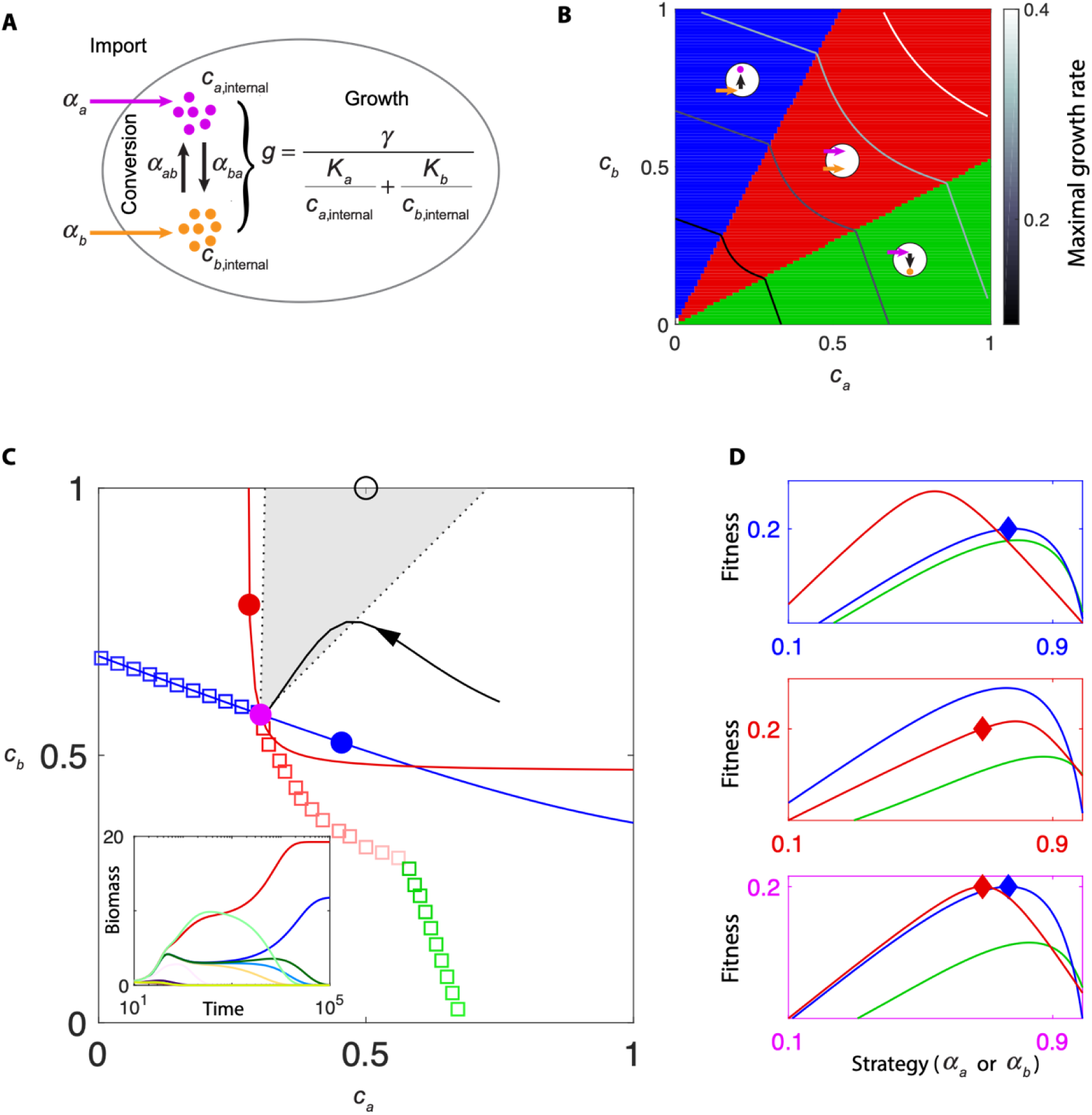
Non-invasible cartels. A. Metabolic model with a trade-off in allocation of internal resources for import of two nutrients plus their interconversion, with both nutrients necessary for growth. B. Three subclasses of maximizing metabolic strategies in chemical space are indicated by background color, and circles with arrows illustrate the metabolic strategies of each subclass. The maximal growth contours for four growth rates (0.1, 0.2, 0.3, 0.4) are marked by gray colors. C. Two maximizing strategies co-creating a non-invasible steady state. At dilution rate 0.2, the maximal growth contour and the corresponding maximizing strategies are shown as colored squares. At a discontinuous point of the growth contour, the supply lines of two distinct metabolic strategies (*Red* and *Blue*) span a gray region, where any supply condition (e.g. black circle) requires the two maximizing strategies to co-create the environment on the discontinuous point. Red and blue dots mark the environments created by species *Red* and species *Blue* alone, and the purple dot marks the environment co-created by *Red* and *Blue*. Black curve with arrows shows a trajectory in chemical space. Inset: competition dynamics of species *Red* and species *Blue* together with 10 other maximizing species with different strategies. D. The fitness landscapes for the three environments in (C) indicated by corresponding box colors. For class Green and Red, the strategy is represented by *α*_*a*_, for class Blue, the strategy is represented by *α*_*b*_.

Following the general protocol described in the previous section, we first identified the maximal growth rates 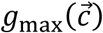 and the corresponding strategy or strategies 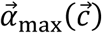 at each point 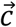 in the chemical space, and generated maximal growth contours for different dilution rates (Fig 5B). The maximal growth contours are not smoothly continuous, nor are the corresponding strategies. In chemical space, three distinct sectors of maximizing strategies appear (Fig 5B, Fig S6A): When nutrient *a* is very low compared to *b*, the maximizing strategy is a “*b-a* converter” which imports *b* and converts it into *a* (blue sector, only *α*_*b*_ and *α*_*ab*_ are non-zero). Symmetrically, when *a* is comparatively high, the optimal strategy is a “*a-b* converter” (green sector, only *α*_*a*_ and *α*_*ba*_ are non-zero). Otherwise, the maximizing strategy is an “importer” which imports both nutrients without conversion (red sector, only *α*_*a*_ and *α*_*b*_ are non-zero). On the border between sectors, the maximal growth contour has a discontinuous slope.

Optimal coexistence occurs at these discontinuous points. If an environment point 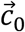 is located in a continuous region of the maximal growth contour, only one maximizing strategy 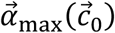 exists for that environment (maximizing strategies along the maximal growth contour are indicated by colored squares in Fig 5C). Supply conditions that make 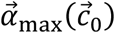 the optimal strategy (i.e. allow 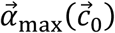 to create the steady-state environment 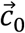) constitute the supply line for 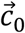 and 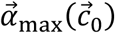. However, at the discontinuous points of the maximal growth contour, where two classes of strategies meet, two different strategies are both maximizing. For example, at the purple dot in Fig 5C a strategy belonging to the “*b-a* converter” class (species *Blue*) and one belonging to the “importer” class (species *Red*) are both maximizing strategies. Each strategy derives a supply line from the purple dot (black dashed line, Fig 5C). The two supply lines span a gray region where no supply line from any single strategy enters. Correspondingly, for any supply conditions inside the gray region, no single species can alone create an environment on the maximal growth contour. For example, under the supply condition shown by the black open circle in the gray region, species *Blue* and species *Red* both create chemical environments that lie within the maximal growth contour (blue and red dots, Fig 5C), and are thus subject to invasion by other species. Nevertheless, the species-specific growth contours of *Blue* and of *Red* intersect at the purple point on the maximal growth contour. Therefore, only when *Blue* and *Red* coexist can they co-create an environment on the maximal growth contour, and thus be resistant to invasion from any other species. Indeed, when we simulate multiple species with different maximizing strategies under the supply condition indicated by the open black circle, species *Blue* and species *Red* are the only two that survive (Fig 5C, inset).

The optimal coexistence of species *Blue* and species *Red* can be understood intuitively from the dynamic fitness landscape. Given a chemical environment, the relation between *α*_*a*_ and growth rate of importer (red curve) or *a-b* converter (green curve), and that between *α*_*b*_ and growth rate of *b-a* converter (green curve) constitute the fitness landscape of species adopting different possible maximizing strategies (Fig 5D). In the environment created by species *Blue* (blue dot in Fig 5C), not only will some importers grow faster than *Blue*, species *Blue* (strategy marked by blue diamond) is not even on the fitness peak of its own class (Fig 5D, upper box). Similarly, in the environment created by species *Red*, the strategy of *Red* is not at the top of the fitness landscape (Fig 5D, middle box). By contrast, in the environment co-created by species *Blue* and *Red* (purple dot in Fig 5C), their strategies are at the top of the fitness landscapes of their own classes and at equal height. For all supply conditions in the gray region, species *Blue* and species *Red* jointly drive the nutrient concentrations to the discontinuous point of the optimal growth contour, and thereby achieve evolutionarily stable coexistence.

### Species creating a new nutrient dimension

One possible solution to the competitive-exclusion paradox is the creation of new nutrient “dimensions” by species secreting metabolites that can be utilized by other species. For example, *E. coli* secretes acetate as a by-product of glucose metabolism. Accumulation of acetate impedes the growth of *E. coli* on glucose (Luli & Strohl, 1990), but the acetate can be utilized as a carbon source, e.g. by mutant strains that emerge in long-term evolution experiments (D’Souza et al., 2018; Rosenzweig, Sharp, Treves, & Adams, 1994).

To explore the possibilities of evolutionarily stable coexistence when species create new nutrients, we used a simplified model to represent multi-step energy generation with a dual-role intermediate metabolite (Fig 6A). A single chemical energy source S is supplied into the chemostat. The pathway for processing S consists of four relevant reactions driven by designated enzymes: External S can be imported and converted into intermediate I_int_ to generate ATP (with a corresponding fraction of the enzyme budget *α*_ATP1_).The intermediate has a dual role in energy production: on the one hand, it positively contributes to ATP production via a downstream reaction (with a fraction of the enzyme budget *α*_ATP2_); on the other hand, it negatively contributes to ATP production through product inhibition of the first energy-producing reaction. To deal with this negative effect of internal intermediate, cells may synthesize transporters (with a fraction of the enzyme budget *α*_exp_) to export intermediate out into environment, where it becomes external intermediate I_ext_. By this reaction, cells can increase the dimension of chemical space from one (S) into two (S and I_ext_). Cells can also import I_ext_ into I_int_ (fraction of enzyme budget *α*_imp_), then use I_int_ as an energy source via the second reaction. (See Methods for details.)

**Figure 6.**
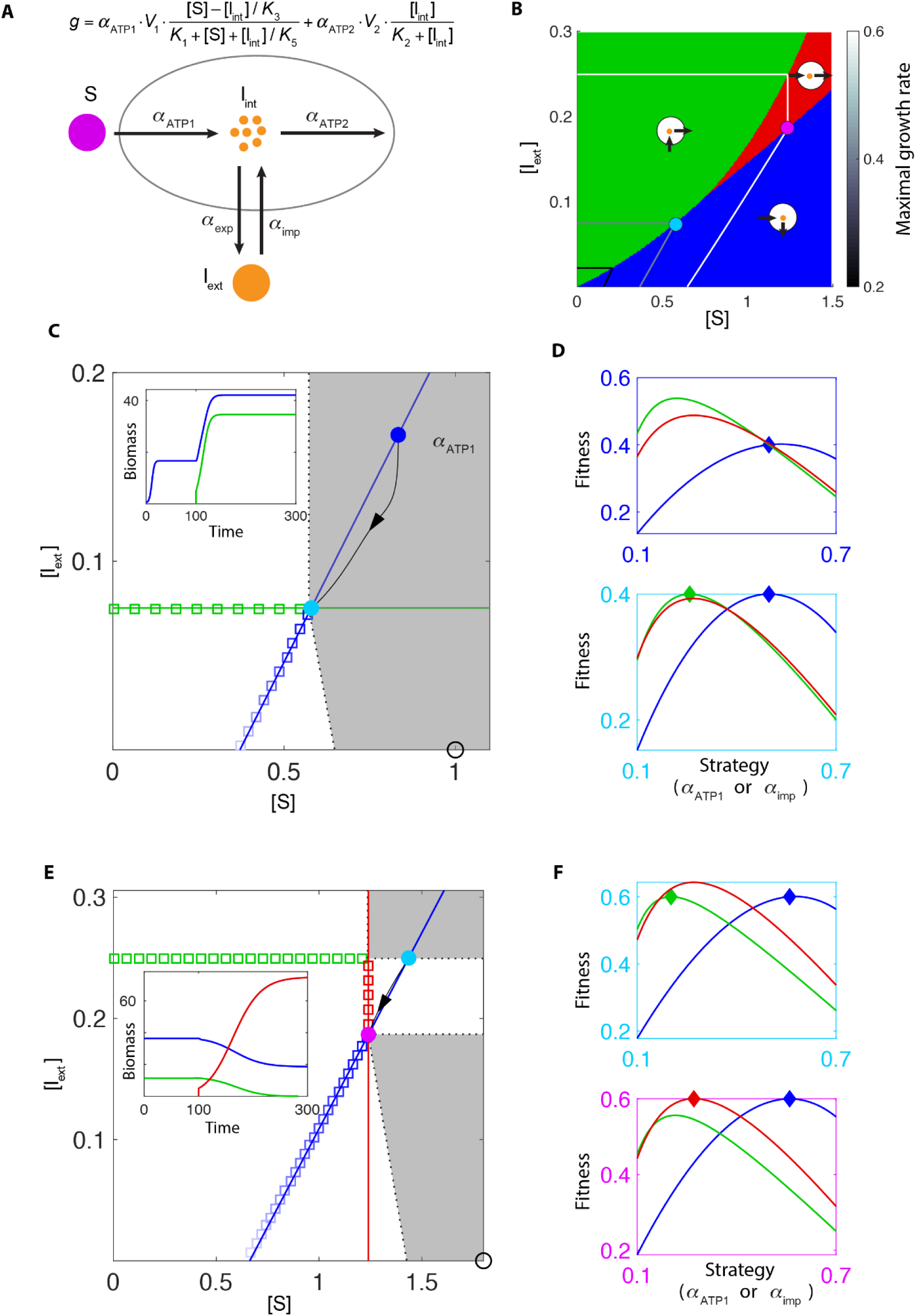
Species creating new nutrient dimensions and achieving evolutionarily stable coexistence. A. Metabolic model with a single supplied nutrient S. Cells allocate enzymes to convert S into internal intermediate I_int_ and produce energy (denoted as “ATP”), export internal intermediate into the chemostat to become I_ext_, import external intermediate, or consume I_int_ to produce ATP. The growth rate is the sum of ATP production (see Methods). B. Three subclasses of maximizing metabolic strategies in chemical space are indicated by background color, and circles with arrows illustrate the metabolic strategies of each subclass. The maximal growth contours for three growth rates (0.2, 0.4, 0.6) are marked by black-to-white colors. C. At dilution rate 0.4, two maximizing strategies co-create a non-invasible environment. The maximal growth contour and the corresponding maximizing strategies are shown as colored squares. At a discontinuous point of the growth contour, the supply lines of two distinct metabolic strategies (*Green* and *Blue*) span a gray region, where any supply condition (e.g. black circle) requires two maximizing strategies to co-create the environment at the discontinuous point. Blue dot marks the environment created by species *Blue* alone, and the cyan dot marks the environment co-created by *Blue* and *Green*. Black curve with arrows shows a trajectory in chemical space. Inset: time course of species biomass, with species *Green* added to the chemostat at time 100. D. The fitness landscapes for two environments in (C) indicated by corresponding colors of the boxes, reflecting the relationship between instantaneous growth rate and resource allocation strategy. For class Blue and Red, the strategy is represented by *α*_ATP1_; for class Green the strategy is represented by *α*_imp_. E. Same as (C), except that the dilution rate is 0.6. Inset: time course of species biomass, starting with *Blue* and *Green*, with species *Red* added to the chemostat at time 100. F. Same as (D), except corresponding to the two steady-state environments shown in (E).

The metabolic strategy 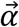 in this model has four components: 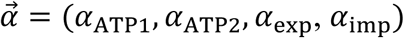. When we examine the maximizing strategies and maximal growth rates in the chemical space, three distinct classes of strategy emerge (Fig 6B). When S is abundant and I_ext_ is low, the maximizing strategies have only two non-zero components, *α*_ATP1_ and *α*_exp_ (Fig S6B), meaning this class of species only imports S, for the first energy-generating reaction, then exports intermediate as waste. Therefore, we call strategies in this class “polluters” (blue section in Fig 6B, Fig S3C). When I_ext_ is high while S is low, the maximizing strategies have two different non-zero components, *α*_ATP2_ and *α*_imp_ (Fig S6B), meaning this class of species relies solely on I_ext_ as its energy source. We call these strategies “cleaners” as they clean up I_ext_ from the environment, (green section in Fig 6B, Fig S6C). When there are comparable amounts of S and I_ext_ present, a third class of maximizing strategies appears: these cells neither export nor import intermediates, but rather allocate all their enzyme budget to *α*_ATP1_ and *α*_ATP2_ to carry out both energy-producing reactions. We call species in this class “generalists” (red section in Fig 6B, Fig S6C).

As shown in Fig 6B, on the borders between classes of strategies in chemical space, the maximal growth contours turn discontinuously. These points of discontinuity, as in the example in the previous section, are chemical environments corresponding to evolutionarily stable coexistence of species from distinct metabolic classes. The classes of optimally coexisting species change with dilution rate. When the dilution rate is low (*d* = 0.4, Fig 6C), at the discontinuous point of the maximal growth contour, the corresponding two maximizing strategies are one polluter (species *Blue*) and one cleaner (species *Green*). Their supply lines span a gray region where both species *Blue* and species *Green* are required to create a steady-state environment on the maximal growth contour. As by assumption we are only supplying the system with S, the supply condition always lies on the *x*-axis of concentration space. For the supply condition shown by the black open circle in Fig 6C, polluter *Blue* creates a chemical environment (blue dot) far from the maximal growth contour. When the cleaner *Green* is added to the system, not only does the biomass of *Blue* increase (inset), but also the steady-state chemical environment moves to the discontinuous point of the maximal growth contour (cyan dot), where both *Blue* and *Green* occupy the peaks of their fitness landscapes (Fig 6D). This result is consistent with the long-term evolution experiment of *E. coli* and also intuitive: polluter *Blue* and cleaner *Green* form a mutually beneficial relationship by, respectively, providing nutrients and cleaning up waste for each other, thereby reaching an optimal cooperative coexistence.

A quite different coexistence occurs at higher dilution rate (*d* = 0.6, Fig 6E). Growth contours at this dilution rate show two turning points, but neither are between the polluter and the cleaner class. One discontinuous point is between the cleaner class (green squares) and the generalist class (red squares), but the gray region spanned by the corresponding supply lines does not cover the *x*-axis and so does not represent an attainable coexistence when only S is supplied. The other discontinuous point is between the generalist class and the polluter class (blue squares). The gray region spanned by the supply lines of the corresponding two maximizing strategies of generalist class (species *Red*) and polluter class (species *Blue*) does cover the *x*-axis. Therefore, a supply condition with only S within the gray region (e.g., the black open circle) leads to the optimal coexistence of generalist *Red* and polluter *Blue* on the discontinuous point (purple dot), despite the fact that they do not directly benefit each other. Indeed, when the generalist *Red* is added to a system with polluter *Blue* and a cleaner *Green*, the cleaner *Green* goes extinct and the biomass of the polluter *Blue* decreases (inset). Nevertheless, the steady-state chemical environment is moved from a cyan dot lying inside the maximal growth contour to the purple dot lying on the maximal growth contour. In the environment of the cyan dot created by cleaner *Green* and polluter *Blue, Blue* is not on the top of the fitness landscape of the polluter class (Fig 6F, upper box). By contrast, for the fitness landscape created by polluter *Blue* and generalist *Red* (Fig 6F, lower box), despite being lower in biomass, *Blue* occupies the top of the landscape. Therefore, the optimal coexistence of this polluter and this generalist does not arise from direct cooperation, but rather from collaborating to defeat other competitors.

## DISCUSSION

Evaluating microbial metabolic strategies within an ecological context is the major focus of this work. Due to the intensity of competition in the microbial world, it is accepted that natural selection has extensively shaped microorganisms’ internal resource allocation strategies and the regulatory mechanisms controlling these strategies (S. Goyal et al., 2010; Liebermeister et al., 2014). Therefore, a quantitative mapping from metabolic strategies to fitness consequences can further our understanding of both regulation and evolution (Bajic & Sanchez, 2019). Many previous studies of metabolic strategies directly assumed the optimization goal of microbial metabolism to be biomass gain, with the chemical environment acting as a fixed input ({Scott, 2014 #86}(Wang & Tang, 2017)) {Schuetz, 2012 #84} (Roller, Stoddard, & Schmidt, 2016){Brophy, 2014 #85}, which simplifies the problem into a search for a maximum on a static “fitness landscape”. However, in the natural world where metabolic strategies compete and evolve, the feedback between species and their environment produces an intrinsically dynamic fitness landscape in which the actions of one species can influence the fitness of all species (Bajic, Vila, Blount, & Sanchez, 2018). One profound example is the Great Oxygenation Event, when cyanobacteria created an oxygen-rich atmosphere (Kasting & Siefert, 2002), causing a massive extinction of anaerobic bacteria but also stimulating an explosion of biodiversity (Schirrmeister, de Vos, Antonelli, & Bagheri, 2013). Therefore, metabolic strategies need to be assessed within an ecological context, taking into consideration not only how species respond to the environment, but also how species construct their own environment.

For the past thirty years, researchers have been utilized various mathematical tools to study flexible fitness landscapes that change with space, time, and population composition.The prerequisite for a intrinscially dynamical fitness landscape – that the species composition influences the fitness of all species in the system – takes a particularly simple form in chemostat-type resource-competition models: Microbes shape their local environment by exchanging metabolites within a shared chemical environment, which determines the growth rate of all cells. Focusing on the chemical environment created by microbial metabolism, we exhibited a set of intuitive and general procedures for analyzing strategies within various metabolic models. Namely, we showed that to compare a set of fixed strategies, the geometric relationships between their growth contours and their steady-state chemical environments yield an immediate prediction for the outcome of competitions. In searching for optimality over the continuous family of strategies, the “maximal growth contour” envelope of all growth contours provides candidates, and the supply condition selects the non-invasible strategy or strategies from among these candidates via the flux-balance curve. Such selection of optimal strategies also supports the conclusion that having the fastest growth rate in an environment does not necessarily imply being the most competitive strategy, as this strategy may shift the environment in an unfavorable direction. To be non-invasible, strategies also need to be able to construct the environment for which they are best suited.

From the perspective of chemostat-related experiment, this work demonstrates the subtleties of controlling nutrient limitation in chemostats. The capacity of species to shape their own environment, even in a system as simple as a chemostat, presents challenges to controlling which nutrient or nutrients are limiting. By traditional definition, if increasing a certain nutrient leads to an increase of a cell’s growth rate, that nutrient is considered “limiting”. However, growth rate is invariant in a chemostat, being set experimentally by the dilution rate, so inferring nutrient limitation requires special attention. For example, if one sees the same cellular responses under different nutrient supplies, what can one conclude? Cells may be creating the same chemical environment out of different supply conditions (*cf*. Fig S1A), or alternatively cells may be transducing different chemical environments into the same physiological response through mechanisms such as “ratio sensing” (Escalante-Chong et al., 2015). Our graphical approach combined with direct measurements of steady-state nutrient concentrations in the chemostat can precisely define and help guide the control of nutrient limitation (Boer et al., 2010). As described above, changes in supply concentrations shift the flux-balance curve, but do not change the shape of the growth contour. Therefore, by experimentally varying the supply conditions and measuring the chemical environment created by cells, the shape of the growth contour can be obtained. The resulting slope of the growth contour provides information on nutrient limitation even in the absence of detailed knowledge about a cell’s metabolism. For example, in the nutrient *a -* nutrient *b* plane, a near-horizontal growth contour indicates *b*-limited growth while a near-vertical growth contour means *a*-limited growth, and an intermediate slope implies that the two nutrients are co-limiting.

Our work also has implications for microbial community assembly. The competitive exclusion principle poses a long-term puzzle in ecology: since species are in constant competition in the natural world, why doesn’t the fittest species outcompete the others and become the sole survivor? Actually, on the basis of simple resource-competition models, it has been argued that the number of stably coexisting species cannot exceed the number of resources, leading to the so-called competitive exclusion principle (Armstrong & McGehee, 1980; Hardin, 1960; Levin, 1970; McGehee & Armstrong, 1977). Nevertheless, tremendous biodiversity manifests in the real world, from environmental surveys to controlled lab experiments (Friedman et al., 2017; Goldford et al., 2018; Maharjan et al., 2006). Our work is not aimed at adding another solution to the paradox of the plankton; rather, our framework helps in summarizing the criteria for coexistence on both ecological and evolutionary timescales. On the ecological scale where a limited number of species with different strategies compete for resources, the intransitivity of competitiveness as a result of fitness landscape deformability is the key to coexistence. Given resource allocation trade-offs, the growth contours of any pair of strategies must intersect, clearly demonstrating why trade-offs prevent a single species from unconditional dominance, allowing various forms of intransitivity under different metabolic models. In the case of substitutable nutrients, the fitness landscape can even be made flat – allowing unlimited coexistence. When the rule of invasion allows non-transitive loops, oscillations and chaos can occur, which have been shown to allow coexistence beyond competitive exclusion (Huisman & Weissing, 1999, 2001). In addition, when multiple nutrients are all essential, the ability of each species to create an environment that favors itself allows for the spontaneous emergence of spatial heterogeneity in an extended system (Fig S5). Meanwhile, on the evolutionary time scale where mutation/adaptation allows searches for the “most suitable” strategies among infinite possibilities, an ongoing threat to diversity is that selection may produce a supreme winner that takes over the habitat. With the dynamic fitness landscape and the maximal growth contour approach, we showed that the condition for evolutionarily stable coexistence is indeed more restricted, occurring only at the discontinuous points of the maximal growth contour. Nevertheless, via the species-environment feedback, a large number of supply conditions can self-organize to the discontinuous point, where multiple species co-create a non-invasible environment where they jointly locate on the peak of the fitness landscape.

Many future directions can follow this work. From the perspective of experiment, our framework can assist in analyzing and interpreting results of microbial evolution in the lab (Van den Bergh, Swings, Fauvart, & Michiels, 2018), where the continual emergence of new mutants under defined experimental conditions suggests an intrinsically dynamic fitness landscape. From the perspective of theory, we do not yet have a rigorous mathematical theorem concerning the conditions for discontinuity of the maximal growth contour, nor proof that discontinuity necessarily leads to evolutionarily stable coexistence. Theoretical developments paralleling those on the general existence of ecologically stable states (De Leenheer et al., 2006; Marsland III, Cui, & Mehta, 2019) would bring a more comprehensive understanding of evolutionarily optimal states in metabolic models. Besides, the metabolic models considered in this work are highly simplified. Going forward, more detailed and experimentally-based models can be examined using the same graphical and dynamical fitness landscape framework.

## METHODS

Programs for this work are coded in MATLAB R2018a. A repository of all tools used to generate the results can be found at: https://github.com/zhiyuanli1987/Qbiotoolbox.git

### Total RNA and total protein measurements

The methods for total RNA and protein measurements shown in Figure S2 are described in Li et al., 2018.

**Supplemental Table 1:**
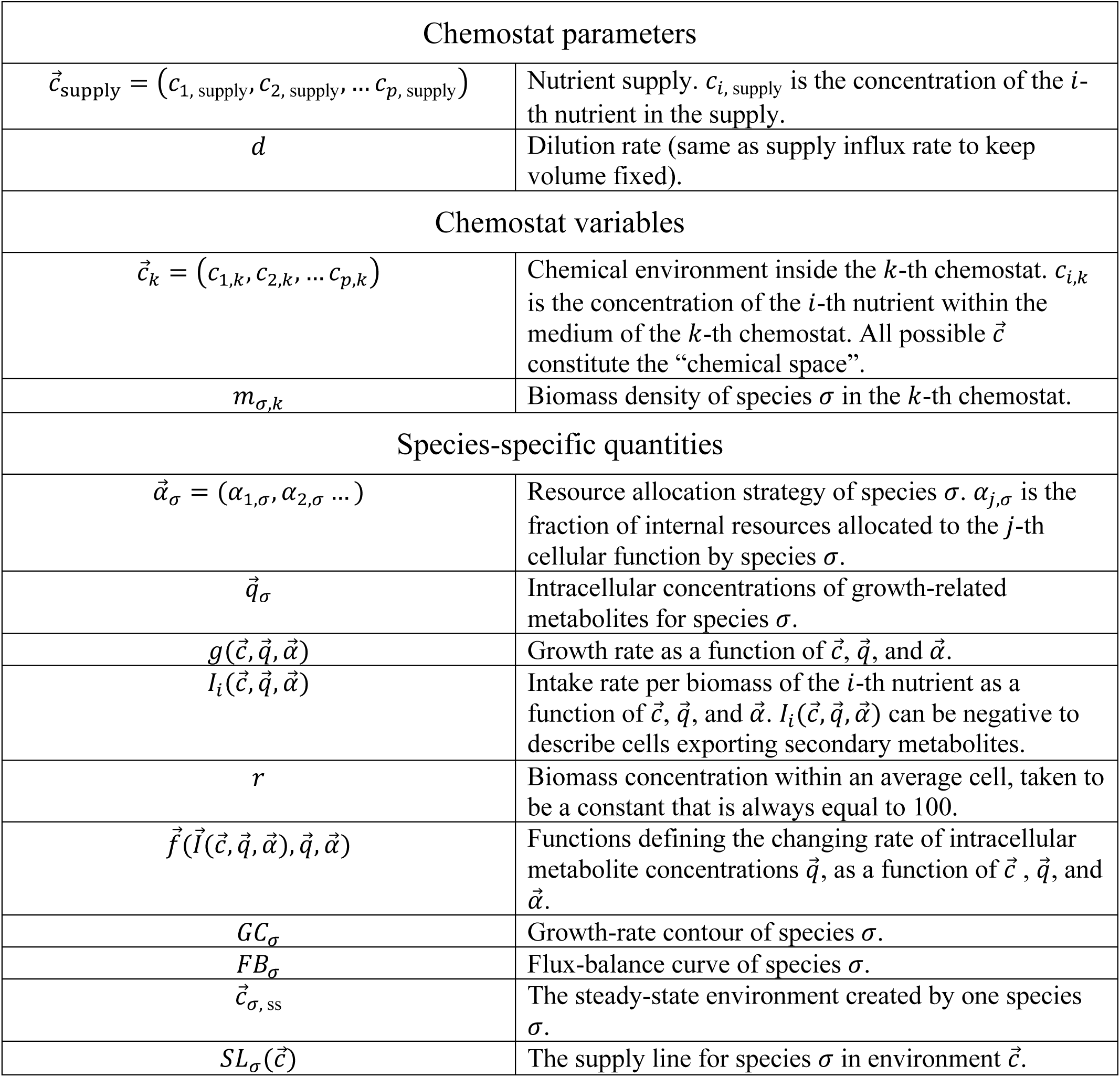
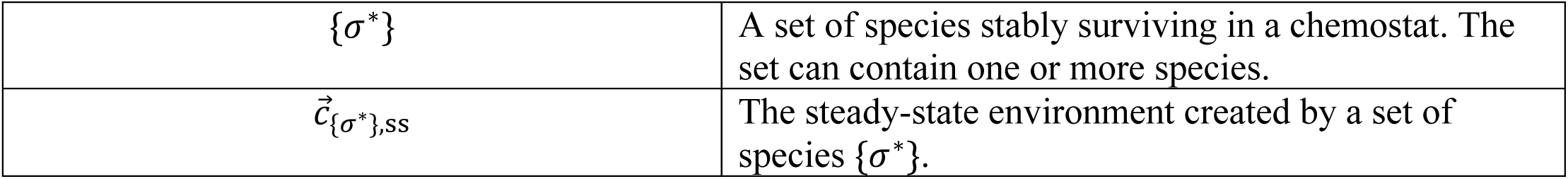
*Symbols*

### Metabolic model and resource allocation strategy

In modeling population dynamics in a chemostat, multiple assumptions need to be made concerning how cells sense the environment, import nutrients, export metabolites, utilize resources, and grow in biomass. Different assumptions result in different metabolic models. Some metabolic models focus on trade-offs in resource allocation, as the amount of internal resources “owned” by a cell, including proteins and energy, is limited. Cells need to allocate these limited resources into different cellular functions, such as metabolism, gene expression, reproduction, motility, maintenance, etc. We use *α*_*j,σ*_ to represent the fraction of internal resources allocated to the *j*-th cellular function of species *σ*, with 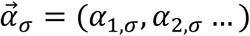 representing the resource allocation strategy of species *σ*. For simplicity, we assume each species has a fixed resource allocation strategy.

### Dynamic equations for a single species in a chemostat

In a chemostat with nutrient supply 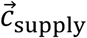, dilution rate *d* and a single species *σ* with fixed strategy 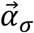 and intracellular metabolite concentration 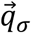, the cell biomass density *m*_*σ*_ and the chemostat nutrient concentrations 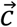 are generally described by the following equations:

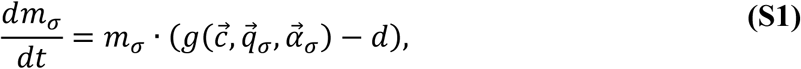

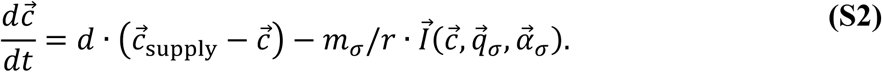

In considering the details of cellular metabolism, one may choose to incorporate the dynamics of intracellular metabolites that originate from nutrient import and influence cell growth. We make the assumption that the biomass concentration *r*, e.g. protein concentration, is constant for cells under all growth conditions. Thus, an increase of total cell mass induces a corresponding increase of total cell volume. *m*_*σ*_ is the cell mass per element of volume in the chemostat, and *r* is the cell mass per element of volume within a cell. For a chemostat-to-cell flux of mass *J*, the concentration of the metabolite in the chemostat decrease by *J*/*V*_chemostat_ while the concentration in cell increase by *J*/*V*_cell_. As a result, the metabolites imported into cells are enriched by a factor of *r*, and metabolites secreted by cells are diluted by 1/*r*. Also, all intracellular metabolites are diluted by cellular growth, which is generally a slow process compared to metabolic reactions and can be ignored in most cases. We use a function 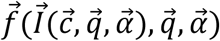 to represent the rate of change of intracellular metabolite concentration 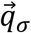:

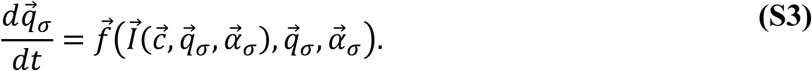

Eq. (S2) represents *p* equations for *p* types of nutrients, and Eq. (S3) represents *h* equations for *h* growth-related intracellular metabolites.

### Species-specific steady state

In the steady state of a chemostat, Eqs. (S1)-(S3) will all be equal to zero.

For intracellular metabolites, as 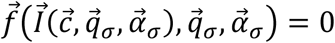 as a result of Eq. (S3) being equal to zero, given an environment 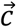, the steady state of 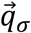 can be expressed as a function of 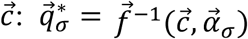.

#### Growth contour (GC)

From the perspective of the environment influencing species, at each constant environment, the steady-state growth rate is fully determined by 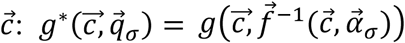. If the biomass of a species is non-zero (*m* ≠ 0), Eq. (S1) requires *g*^*^ = *d*. In the *p*-dimensional chemical space, this requirement defines a (*p* − 1)-dimensional surface, constituted by all environments 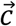 that support an equal-to-dilution growth rate. This surface reduces to the zero-growth isoclines in contemporary niche theory when the chemical space is two-dimensional and the growth rate *g* solely relies on 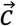 monotonically, but is not necessarily limited by the nutrient dimension or the form of the growth function. For convenience, we name this surface the “growth contour” (GC) for species *σ*:

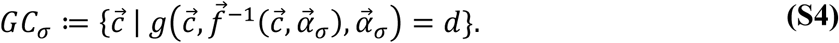

An example of growth contours is shown in Fig 1D.

#### Flux-balance curve (FB)

Eq. (S2) describes how species act on the environment. In steady state, the influx, out-flux, and consumption by species should be balanced for each nutrient, which enables calculation of the biomass density-to-dilution ratio for every 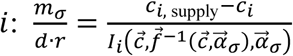. For a *p*-dimensional chemical space, there are *p* equations for the same value of 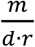. This leads to a one-dimensional curve in the chemical space, which we name the “flux-balance curve” (FB), defined as:

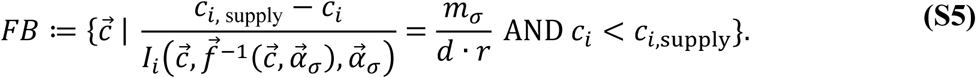

For example, for two nutrients *a* and *b*, the flux balance curve is: 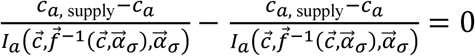, as demonstrated in Fig 1B and Figure S1.

In chemical space, the steady-state environment 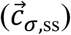 with non-zero biomass of species *σ* will be located at the intersection of the growth contour and the flux-balance curve. This environment is constructed by the species *σ* via its consumption of nutrients. If 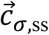 exists, this species can survive in the chemostat. Otherwise, this species will be washed out by dilution even without competition from other species. For the following discussion, we only consider species that can survive when alone in a chemostat.

#### Supply line (SL)

The flux-balance curve is determined by the supply condition 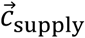. In many cases, it is helpful to derive the supply conditions that enable a species *σ* to construct a steady-state environment 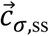. All possible values of 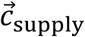that can produce a given 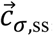, form a straight line in the space of supply concentrations, described by:

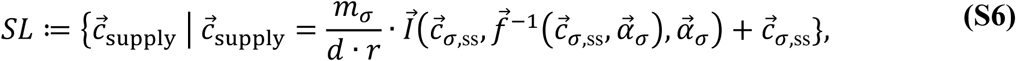

with varying non-negative values of *m*_*σ*_ /*d*. An example of a supply line is shown in Fig S1A.

### Dynamic equations for multiple species in a chemostat

In nutrient competition models, multiple species (*σ* = 1 … *n*) each with biomass density *m*_*σ*_ compete for resources. They have species-specific growth rates 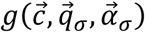 and import rates 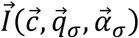, yet all experience the same chemical environment 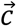. Therefore, Eq. (S1) and Eq. (S3) remain the same for each species, while the rate of change of chemostat nutrient concentrations is influenced by the summed action of all species:

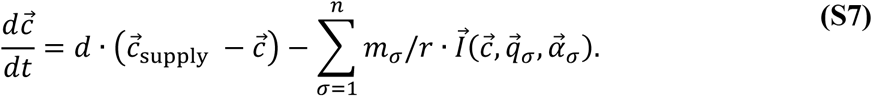

### Multiple species steady state

Multiple species, even if each alone can survive in chemostat, do not generally coexist when competing together. For a system starting with *n* different species, the stable steady state contains *n*^*^ (1 ≤ *n*^*^ ≤ *n*) species with non-zero biomass. We define these *n*^*^ surviving species as a stable species set {*σ*^*^}, and mark the steady-state environment created by this set as 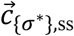. If *n* > 1, according to Eq. (S1), 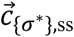 must be located at the common intersection of growth contours formed by every species in {*σ*^*^}.

### Invasion

Invasion is defined as introducing a small number of invaders (with biomass density *m*_inv_) to a steady-state chemostat occupied by a set of local species. At the time of introduction, if the invader can increase in biomass 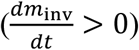, the invasion is successful; otherwise if the invader decreases in biomass 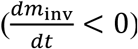, the invasion is unsuccessful. If the biomass stays constant 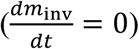, the species is neutral with respect to the local species.

In evaluating invasion by a species *σ* with strategy 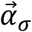 of any environment 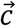, we make two assumptions:

1. The biomass of the invader is so small that it does not disturb the environment at the time of introduction.

2. There is a separation of timescales such that the concentrations of intracellular metabolites reach equilibrium instantaneously at the time of introduction of the invader, therefore Eq. (S3) is always equal to zero and 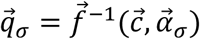 holds.

Therefore, we define the “invasion growth rate” of a species *σ* with strategy 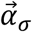 introduced into environment 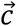 as:

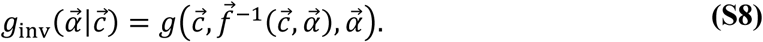

#### Invasion zone

By definition, the growth contour of the invader *GC*_inv_ divides the chemical space into two regions: an “invasion zone” that includes all environments where the invader has an invasion growth rate higher than dilution, and “no-invasion zone” where the invader has an invasion growth rate lower than dilution. If the steady-state environment constructed by local species 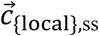 is located within the invasion zone of the invader, 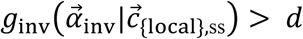, therefore 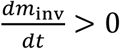 by Eq. (S1), and the invasion is successful; otherwise, if 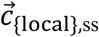 is located outside of the invasion zone of the invader, 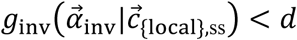, and the invasion is unsuccessful. If 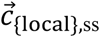 locate exactly on the growth contour, it is neutral.

Two examples of this rule of invasion are presented in Fig 1C and 1D.

If the growth rate monotonically increases with the concentration of each nutrient, it can be proven that the invasion zone is always above the growth contour of the invader (an environment 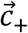 “above” the growth contour *GC*_inv_ is defined as 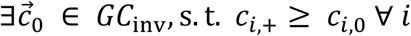. If the growth rate is not monotonically increasing with nutrient concentrations, identifying the invasion zone requires more model-specific analysis.

### Fitness landscape

We quantified the fitness landscapes in the chemostat via the relationship between metabolic strategies 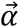 and the invasion growth rates of an invader adopting strategy 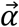 in a given chemical environment 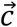. Specifically,

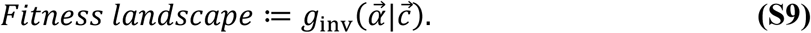

Each environment 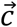 defines a fitness landscape. A set of species {*σ*^*^} constructs a steady-state environment 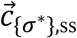 and a corresponding fitness landscape 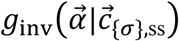. Some examples of fitness landscapes are shown in Figs 2C, 3C, 4B-C, 5D and 6D-F.

### Non-invasible /optimal/ evolutionarily stable strategies

A set of species {*σ*^*^}_opt_ is non-invasible, aka optimal or evolutionarily stable, if no other species can invade the steady-state environment constructed by {*σ*^*^}_opt_:

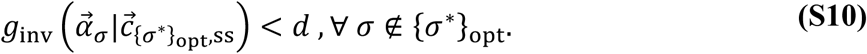

Equivalently, Eq. (S10) can be expressed as “a set of species {*σ*^*^}_opt_ construct a fitness landscape which places themselves on the top”, according to Eq. (S9).

The steady state constructed by {*σ*^*^} is influenced by the supply 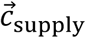 and the dilution rate *d*. For different chemostat parameters, the non-invasible species set {*σ*^*^}_opt_ may be different. In the following steps, we described a generally applicable protocol for obtaining the non-invasible strategies:

#### 1. Maximal growth rates and maximizing strategies

In a metabolic model with trade-offs in resource allocation, the maximizing resource allocation strategy 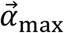 under a given environment 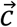 is defined as the strategy that maximizes invasion growth rate:

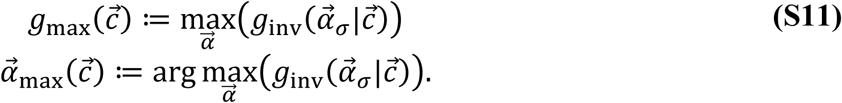

#### 2. Maximal growth contour

For a given dilution rate *d*, all environments that support a maximal growth rate of *d* constitute the “maximal growth contour”:

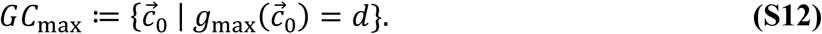

*GC*_max_ is generally formed by many species, with each species adopting the maximizing strategies 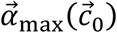 corresponding to one environment 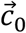 on the maximal growth contour. *GC*_max_ is outside of the invasion zone for any species *σ*. (Otherwise, if a species *σ* could invade an environment 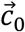 on 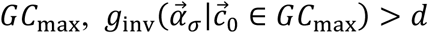, this would directly violate the requirement by Eqs. (S11) and (S12) that 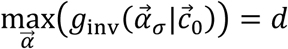 Therefore, the necessary and sufficient condition for a set of species to be evolutionarily stable, is to construct a steady-state environment on the maximal growth contour:

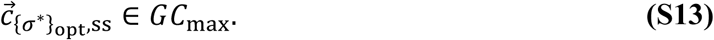

Therefore, a strategy belonging to the non-invasible set must be a maximizing strategy. An example of maximal growth contour is shown in Figs 4D, 5B-C, and 6D

#### 3. Non-invasible strategy

Nevertheless, adopting one of the maximizing strategies along the maximal growth contour does not guarantee that a species will satisfy Eq. (S13) and become non-invasible, as a maximizing strategy for environment 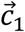 may end up constructing a different environment 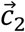. To identify a non-invasible species for supply condition 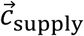, the flux-balance condition needs to be considered, with the strategies maximized at each environment. This requirement forms a “maximal flux-balance curve” in the chemical space:

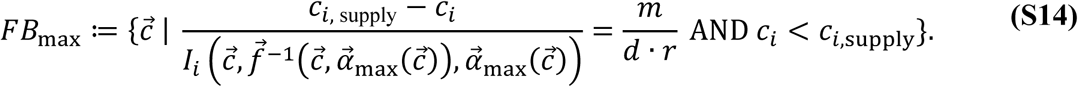

If the intersection of the maximal growth contour and the maximal flux-balance curve exists, it is the evolutionarily stable environment under dilution rate *d* and supply condition 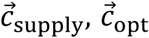.The maximizing strategy for this environment, 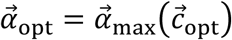, constructs the environment 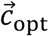, and is evolutionarily stable.

#### 4. Evolutionarily stable coexistence at the discontinuous points of the maximal growth contour

Inversely, for each environment 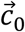 on the maximal growth contour, all supply conditions that enable the maximizing strategy of 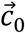 to become the non-invasible strategy can be calculated from the supply line according to Eq. (S6):

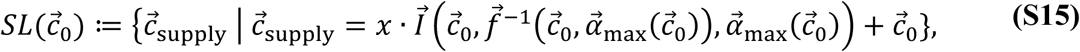

for any non-zero value of *x*. Some examples are shown in Fig 4D.

When there are discontinuous points on the maximal growth contour, there can be “gaps” in the nutrient supply space, where no single strategy on the maximal growth contour satisfies Eq. (S14). Under this condition, more than one strategy is required to co-create an environment on a discontinuous point of the maximal growth contour. Therefore,_discontinuous points of the maximal growth contour permit evolutionarily stable coexistence, where {*σ*^*^}_opt_ contains more than one species. Two examples of such discontinuities and coexistence are shown in Fig 5 and Fig 6.

### Metabolic models

Different assumptions can be made regarding the metabolic models 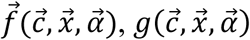, and 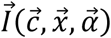, focusing on various aspects of cellular growth. Different assumptions lead to distinct classes of metabolic models with various results. Nevertheless, our analysis schemes, including the invasion geometry, fitness landscape, and evolutionary stable strategies, are generally applicable for various metabolic models. In this work, we used five metabolic models to illustrate multiple aspects of the species-environment feedback:

#### 1. Metabolic model with two essential nutrients

When two nutrients are both essential for growth, such as nitrogen and phosphorus, and both require a substantial allocation of resources for import, the system can be abstractly modeled as shown in Fig 4A. In this metabolic model, we assume an exact trade-off between the allocation of limited resources to import nutrient *a* or nutrient *b*. The fraction of resources allocated to import nutrient *a* is represented by *α*_*a*_, thus leaving a fraction *α*_*b*_ = 1 − *α*_*a*_ to import nutrient *b*. The import rate of nutrient *i* is assumed to follow the Monod equation as a function of nutrient concentration, and is proportional to *α*_*i*_ :

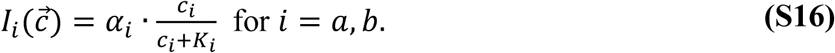

Import of both nutrients is required for cell growth:

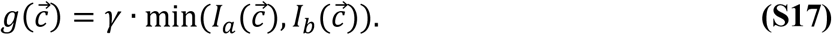

For this model, for simplicity we do not explicitly consider intracellular metabolites. Rather, import directly determines growth. In this model, a “species” is defined by its value of *α*_*a*_. Nutrient limitation can be clearly quantified in this system: if 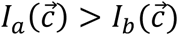, the system is limited by nutrient *b*; if 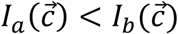, the system is limited by nutrient *a*.

A species with the following parameters was used to generate Fig S1, focusing on how supply conditions and dilution rate influence nutrient limitation:

| $K_a$ | $K_b$ | $\gamma$ | $\alpha_a$ |
| --- | --- | --- | --- |
| 0.7 | 1.3 | 10 | 0.3 |

In Fig S1A, to demonstrate how species construct the same environment out of different supply conditions, the chemostat dilution rate was set to *d* = 1, and three supply conditions were used: 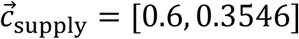 (purple), 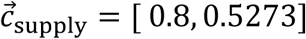(cyan), and 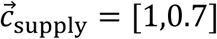 (blue). In Fig S1B, to demonstrate how dilution rates may switch the limiting nutrient, we used the same supply condition as the blue condition in Fig 1D 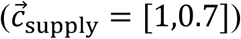, and three dilution rates: 0.5 (yellow), 1 (red), and 1.6 (deep red).

Species with following parameters were used to generate Fig 4B-D:

| $K_a$ | $K_b$ | $\gamma$ |
| --- | --- | --- |
| 0.5 | 0.5 | 10 |

The strategy *α*_*a*_ varies for different species. In Fig 4B, Species *Blue* has *α*_*a*_ = 0.35, species *Red* has *α*_*a*_ = 0.65. In Fig 4C, we started with species *Blue* and species *Red*. We then generated the fitness landscape for each species at the steady-state environment it constructed, then chose the strategy that maximized invasion growth rate for this fitness landscape to generate a new species, and iterated this process five times. The species *Black* has *α*_*a*_ = 0.5.

In generating Fig 4D, we followed the protocols described in section “Non-invasible / evolutionarily stable strategies”.

#### 2. Metabolic model with substitutable nutrients

When two nutrients are mutually substitutable for growth, such as glucose and galactose, the system can be described by metabolic model as shown in Fig 2A. The trade-off and import functions are taken to be the same as in Model 1: *<u>metabolic model with two essential nutrients</u>*. However, import of the two nutrients contributes additively toward growth rate:

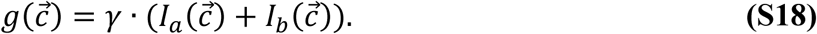

A species is defined by its value of *α*_*a*_.

For this model, all growth contours intersect at one point. The growth contour of species *σ* satisfies the equation: 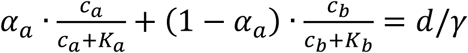. Regardless of the value of *α*, the environment 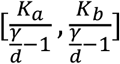 is always on the growth contour.

Species with the following parameters were used to generate Fig 1C-D and Figure 2.:

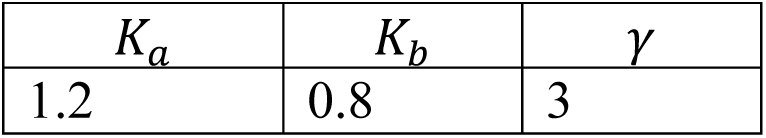

The strategy *α*_*a*_ varies for different species. In Fig 1C-D and Fig 2B-C, Species *Blue* has *α*_*a*_ = 0.2, species *Red* has *α*_*a*_ = 0.6. Supply conditions are different among the three figures: in Fig 1C, 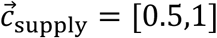; in Fig 1D, 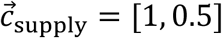; in Fig 2B, 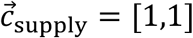.

All conditions in Fig 2D are the same as in Fig 2B other than that five additional species are added to the system. Their strategies are indicated by the legend at the right.

#### 3. Metabolic model with substitutable nutrients that require assimilation

In cells, the assimilation of imported raw material, such sugars, into biomass such as proteins, takes multiple steps and enzymes and consumes a considerable amount of energy. When the resources allocated to nutrient assimilation are considered, a cell’s strategy becomes more complex. A mathematical model involving three substitutable nutrients *a, b, c* that need assimilation is shown in Fig 3A, with *α*_*i*1_ represents the fraction of resources allocated to importing nutrient *i* into internal metabolite, and *α*_*i*2_ represents the fraction of resources allocated to assimilate the internal *i* into biomass. In this model, the import rate has a similar form to the previous two models,

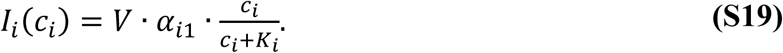

The internal metabolite concentration *c*_*i*,internal_ has an influx of *r* · *I*_*i*_ (*c*_*i*_), meanwhile, it is diluted by cell growth at the rate *g*. We assume all nutrients are substitutable therefore the internal pools contribute via summation to growth, and are converted into biomass at a rate *k* · *α*_*i*2_ · *c*_*i*,internal_:

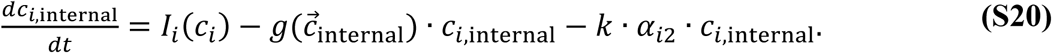

Therefore, the mass converted into biomass per unit time per unit volume is: ∑_*i*_(*k* · *α*_*i*2_ · *c*_*i*,internal_), and the growth rate defined as the relative gain of total biomass *M* is:

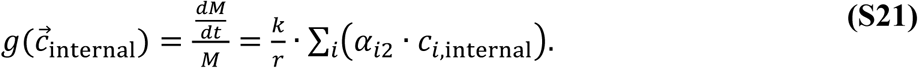

In generating Fig 3B-C, the chemostat parameters were: 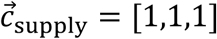, and *d* = 1, and the species parameters were:

| $V$ | $K_i (i = a, b, c)$ | $k$ |
| --- | --- | --- |
| 1000 | 0.5 | 11 |

The three species allocate their resources differently:

| Strategies | $\alpha_{a1}$ | $\alpha_{a2}$ | $\alpha_{b1}$ | $\alpha_{b2}$ | $\alpha_{c1}$ | $\alpha_{c2}$ |
| --- | --- | --- | --- | --- | --- | --- |
| *Red* | 0.15 | 0.2 | 0.1 | 0.25 | 0.26 | 0.04 |
| *Green* | 0.26 | 0.04 | 0.15 | 0.2 | 0.1 | 0.25 |
| *Blue* | 0.1 | 0.25 | 0.26 | 0.04 | 0.15 | 0.2 |

To generate the Fig S3, all other parameters are the same, other than *k* = 10.

#### 4. Metabolic model with essential nutrients that can be interconverted

If two nutrients are both essential for growth, and a cell is able to convert one nutrient into another albeit at a certain cost, as shown in Fig 5A, metabolic trade-offs involve the following four elements of the allocation strategy 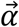 :

*α*_*a*_: Fraction of resources allocated to import nutrient *a*.

*α*_*b*_: Fraction of resources allocated to import nutrient *b*.

*α*_*ab*_: Fraction of resources allocated to convert internal *b* into *a*.

*α*_*ba*_: Fraction of resources allocated to convert internal *a* into *b*.

To implement trade-offs, the sum of elements of 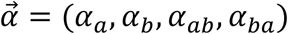 is taken to be equal to 1. In this metabolic model, cells internalize nutrient *a* and nutrient *b* from the chemostat to supply internal concentration of nutrients, *c*_*a*,internal_ and *c*_*b*,internal_. Meanwhile, the internal nutrients can be converted into each other. Nutrients also diffuse in and out of the cell passively with rate *β*. Cell growth requires both internal nutrients, and depletes them in a fixed proportion. In this model, the growth rate of a cell is taken to be:

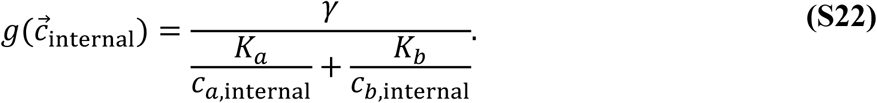

The net import rate, including passive diffusion, is:

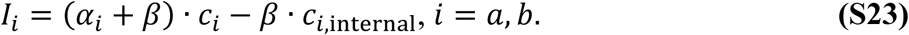

Therefore, the dynamical equations for the internal nutrients are:

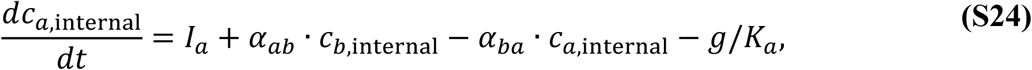

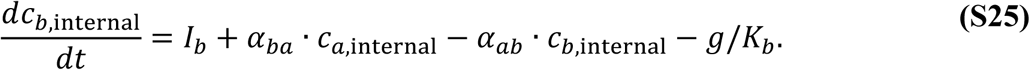

A species is defined by its value of 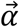.

This metabolic model was used to demonstrate how to obtain locally optimal strategies and cartels, as shown in Fig 5. The parameter values used to generate the plots in Fig 5B-D were:

| $\gamma$ | $K_i (i = a, b)$ | $\beta$ |
| --- | --- | --- |
| 1 | 1 | 0.2 |

In generating Fig 5B, we searched for the maximizing strategies in the chemical space, and classified them by their non-zero values. Maximal growth contours for four dilution rates: 0.1, 0.2, 0.3, 0.4 are shown from black to gray and white colors.

In generating Fig 5C, the chemostat parameters were set to 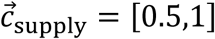, and *d* = 0.2. The maximal growth contours for *d* = 0.2 were drawn, along with maximizing strategies along the contour shown as squares with colors corresponding to their sub-classes. At the discontinuous point of the maximal growth contour where the “converter” and the “importer” converge, the distinct two maximizing strategies are denoted species *Red* and species *Blue*. In generating the competition dynamics in inset, additional to the species *Red* and species *Blue*, ten other maximizing strategies along the maximal growth contours were chosen.

#### 5. Metabolic model with multiple energy generating steps

Cell growth is also tightly coupled with energy production. For example, with a single carbon supply as the energy source, cells employ multi-step reactions to generate multiple ATP molecules. Each step requires dedicated enzymes. The reaction intermediates, such as acetate, usually have dual roles: on the one hand, they positively contribute to ATP production via downstream reactions; on the other hand, they negatively contribute to ATP production by hampering upstream reactions. To deal with the negative effects of intermediates, cells may transport them out into the environment, generally with some metabolic cost for transporters. On the other hand, cells can also uptake such intermediates and use them as an energy source. We abstract such a process by the model shown in Fig 6A. A single chemical energy source S is supplied into the chemostat, which can be converted into intermediate I by cells. Four reactions are possible in this model, each mediated by a specific enzyme:

1. Import the resource S into the cell and convert it into internal intermediate I_int_ to extract energy (e.g. ATP). The fraction of the model enzyme budget allocated to this reaction is *α*_ATP1_. We assume the reaction is reversible, with the concentration of S contributing positively to the reaction rate while the concentration of I_int_ contributes negatively:

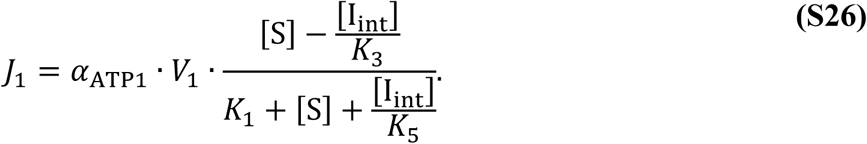

2. Process I_int_ via a downstream reaction to obtain more energy. The fraction of enzymes being allocated to this reaction is *α*_ATP2_. For this model system, it does not qualitatively influence the final results whether this reaction is product inhibited, so we neglect product inhibition. For simplicity, we assume this reaction has Michaelis–Menten form:

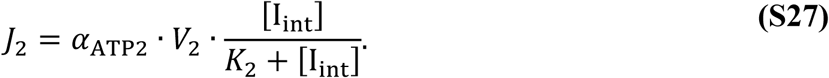

3. Export the internal intermediate out into the environment by diffusion, with a fraction of proteins *α*_exp_ allocated to channels that allow the excretion of the intermediate into the environment to become external intermediate I_ext._

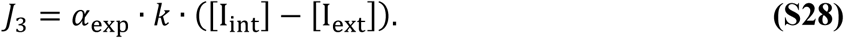

4. Import the external intermediate into cells, with a fraction of proteins *α*_imp_ allocated to the import process. To reflect the property of the internal intermediate in inhibiting this transport reaction, the rate for this process is also product-inhibited:

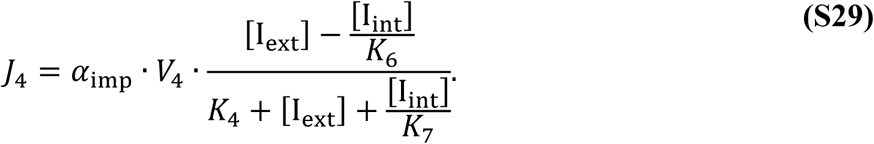

Under this model, the rate of change of the concentration of the energy source S in the chemostat is:

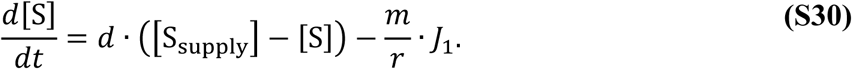

The rate of change of the external intermediate concentration in the chemostat is:

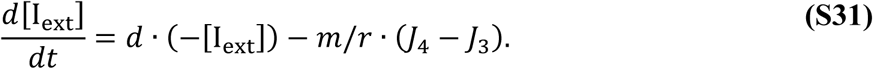

The concentration of the intracellular metabolite I_int_ follows the equation:

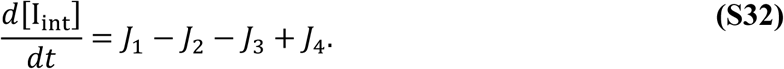

The growth rate is a weighted sum of the ATP produced by *J*_1_ and *J*_2_:

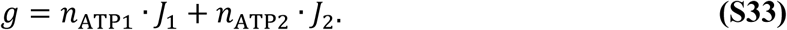

In generating plots in Fig 6B-F, the species parameters were:

| $V_1$ | $V_2$ | $k$ | $V_4$ | $K_1$ | $K_2$ | $K_3$ | $K_4$ | $K_5$ | $K_6$ | $K_7$ | $n_{\text{ATP1}}$ | $n_{\text{ATP2}}$ |
| --- | --- | --- | --- | --- | --- | --- | --- | --- | --- | --- | --- | --- |
| 5 | 1 | 8 | 10 | 0.5 | 0.5 | 0.5 | 0.1 | 0.5 | 15 | 10 | 1 | 1 |

Maximal growth contours for dilution rates 0.2, 0.4, and 0.6 are shown in Fig 6B.

For Fig 6C-D, the chemostat parameters are: *S*_supply_ = 1, *d* = 0.4.

For Fig 6E-F, the chemostat parameters are: *S*_supply_ = 1.8, *d* = 0.6.

A summary of maximizing strategies in chemical space is shown in Fig S4.

### Dynamic equations for multiple species in a chain of chemostats

Real ecosystems seldom exist in isolation. We modeled interconnected ecosystems via a chain of chemostats labeled *k* = 1 to *k*_tot_ (Fig S5A). Each chemostat exchanges medium and cells at leakage rate *l* with its two neighboring chemostats (if *k* = 1 or *k* = *k*_tot_, there is only one neighbor). The chemostat parameters 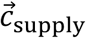and *d* are taken to be identical for all chemostats.

For the *k*-th chemostat, the dynamical equations for the biomass density of species *σ* and the concentration of the *i*-th nutrient are:

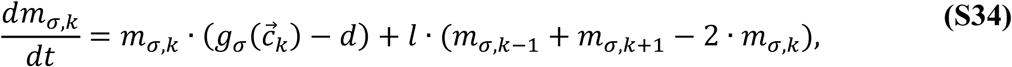

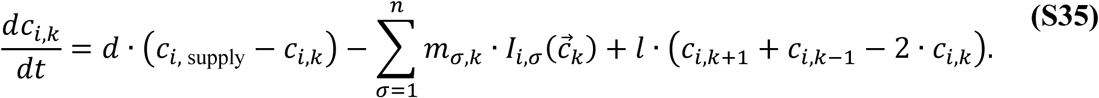

A steady-state solution to these equations is shown in Fig S5, using the same growth and import models and parameters as in Fig 4, with the leakage rate set to be *l* = 1.

## ACKNOWLEDGEMENTS

We thank Simon Levin for insightful discussions. Zhiyuan Li was supported by the Princeton Center for Theoretical Science and the Center for the Physics of Biological Function. This work was supported by the National Institutes of Health Grant R01GM082938 and by the National Science Foundation, through the Center for the Physics of Biological Function (PHY-1734030).

## SUPPLEMENTAL FIGURES

**Figure S1.**
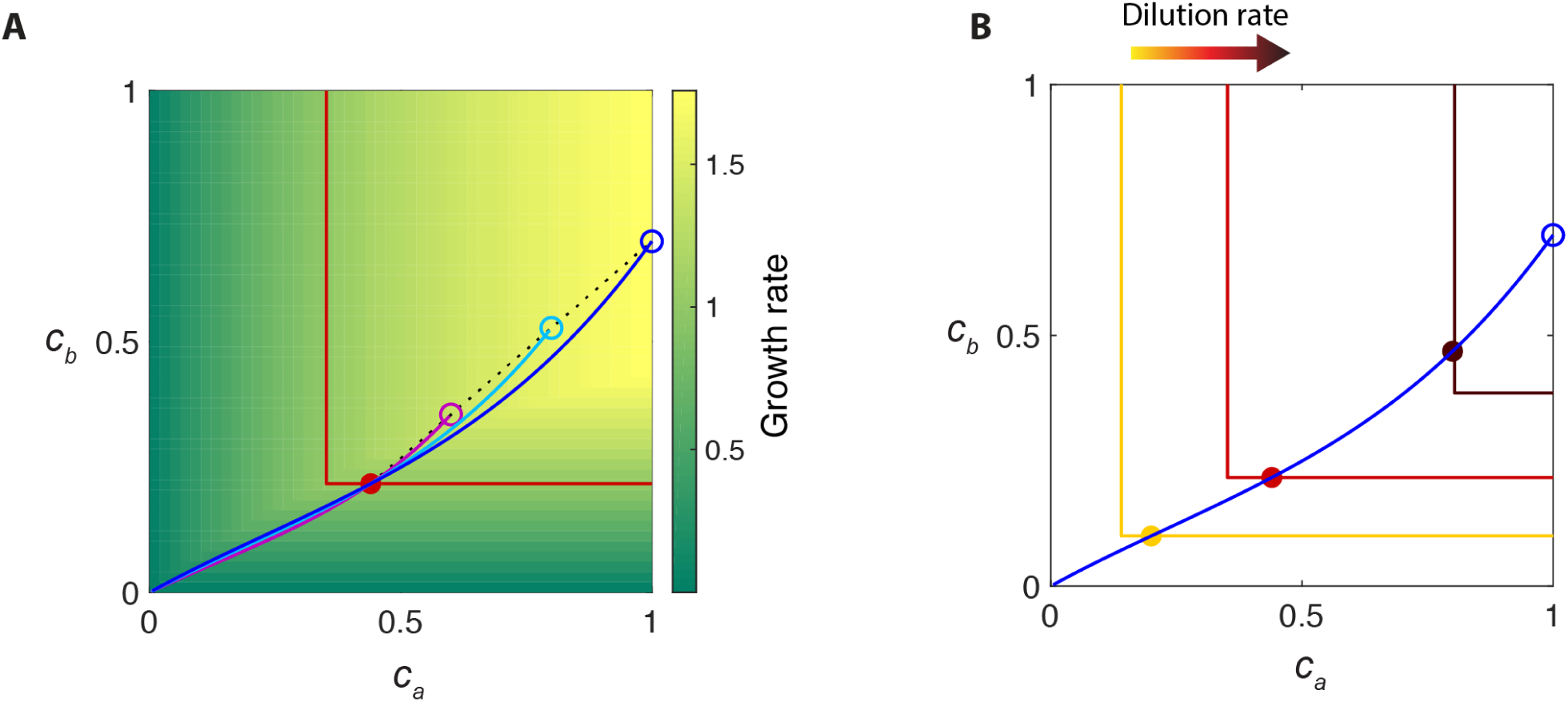
How supply concentrations and dilution rate separately influence the shapes of nullclines and the steady-state environment. A. Various supply concentrations can lead to the same steady-state chemical environment. Background color indicates the growth rate of cells as a function of nutrient concentrations *c*_*a*_ and *c*_*b*_, with the growth contour shown by the red curve. The supply line for the steady-state environment (purple dot) is shown as a dotted black line. Different supply concentrations (*c*_*a,*supply_ and *c*_*b,*supply_) along the supply line are marked by purple, cyan, and blue circles, with the corresponding flux-balance curves shown in the same colors. B. Dilution rate can flip nutrient limitation. The external supply condition is marked by a blue circle, and the flux-balance curve for this supply is shown in the same color. Three growth contours with increasing dilution rates are shown from yellow to deep red, and the corresponding steady-state environments are shown in colored dots.

**Figure S2.**
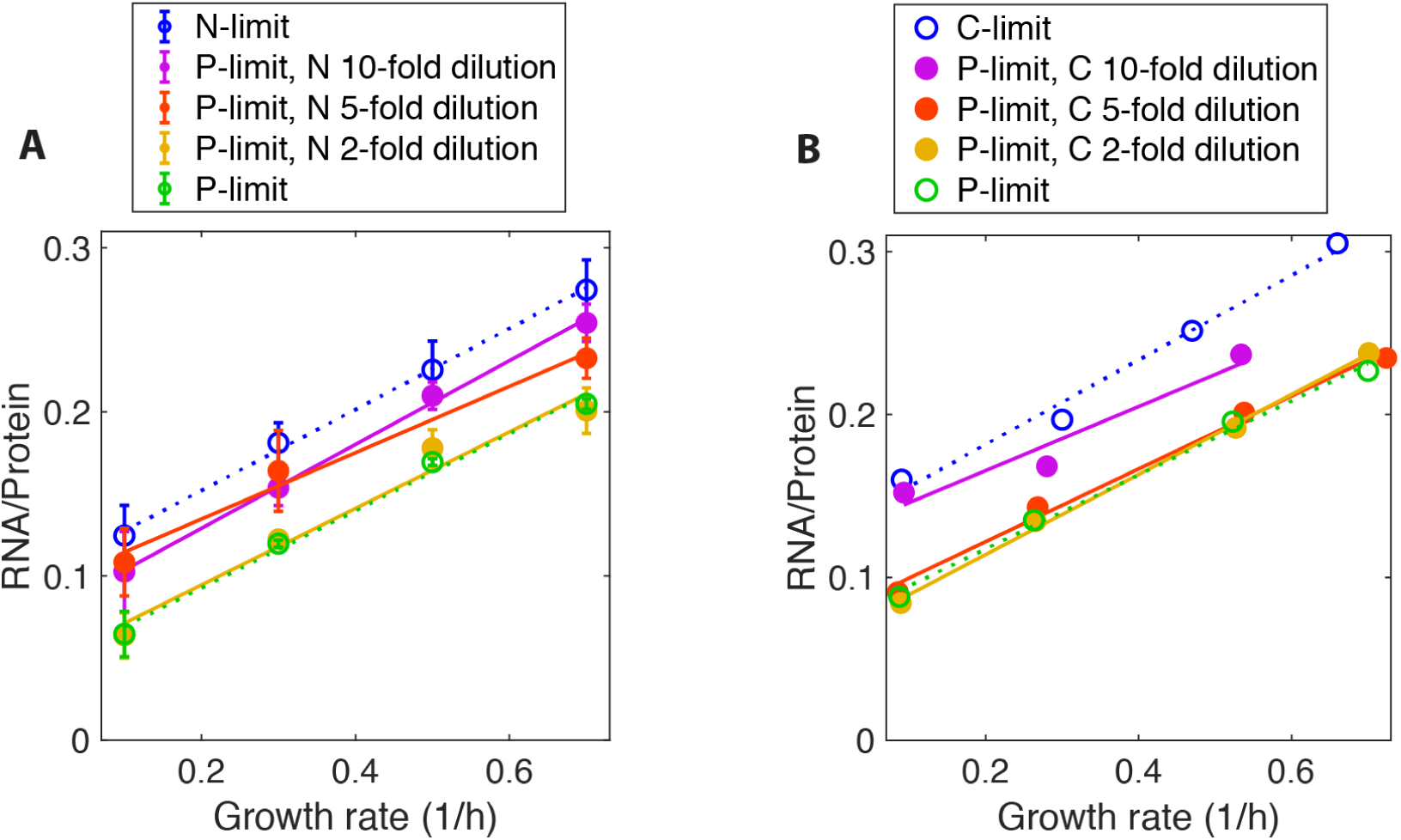
Nutrient supply shifts the relationship between RNA/Protein ratio and growth rate in chemostat. A. The relationship between ribosome abundance represented by RNA/Protein ratio (*y*-axis) and growth rate (*x*-axis) of *E. coli* cultured in chemostats from phosphorus limitation (P-limited, green open circles and dotted line) to nitrogen limitation (N-limited, blue open circles and dotted line). Starting from the P-limited condition, data for decreasing the supply concentration of nitrogen by 2, 5, and 10-fold are shown as solid dots and corresponding best-fit lines. Each measurement was repeated three times and standard errors are shown by bars. C. Same as (B), but for phosphorus and carbon limitation instead of phosphorus and nitrogen limitation. Starting from the P-limited condition, data for decreasing the supply concentration of carbon by 2, 5, and 10-fold are shown as solid dots and corresponding best-fit lines.

**Figure S3.**
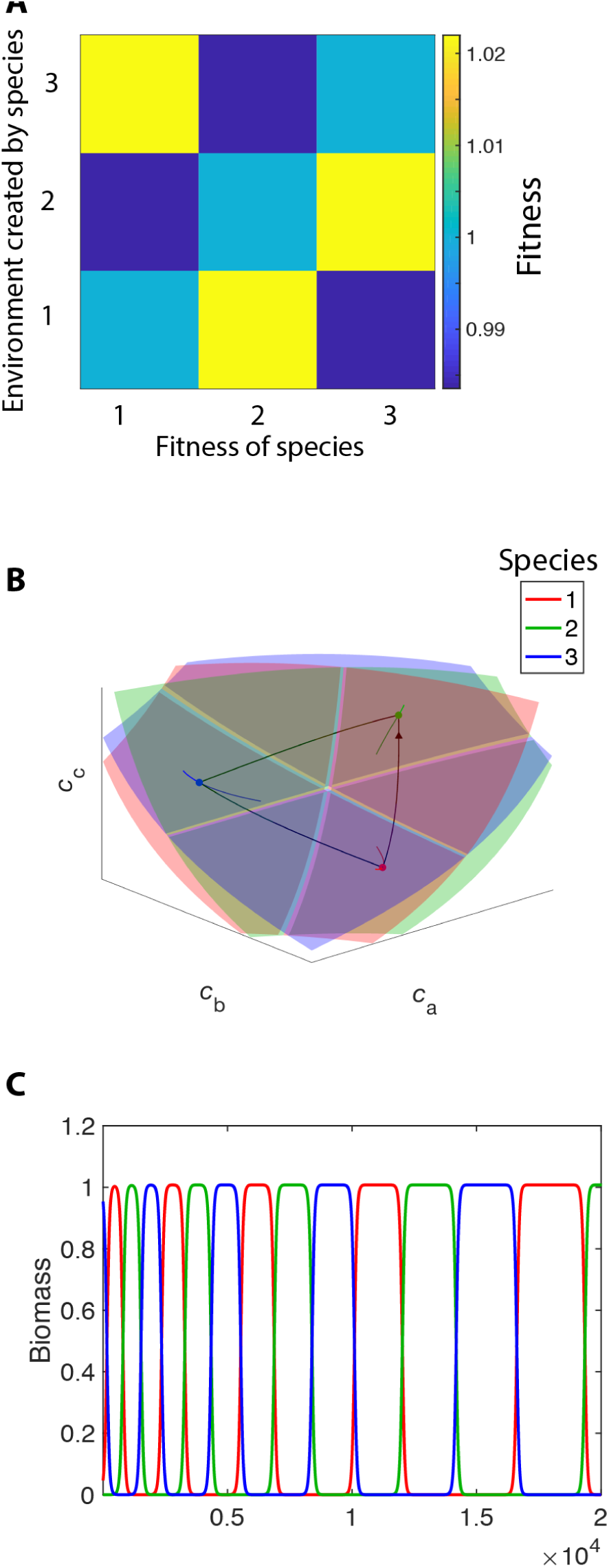
Rock-paper-scissor fitness landscape and heteroclinic cycle. A. The fitness of Species 1, 2, and 3 in the steady-state environment constructed by species 1, 2, and 3 for the model in Fig 3. B. Growth contours (surfaces), flux-balance curves (lines), and steady-state nutrient concentrations (dots) for three species in a three-dimensional nutrient space, with a different conversion speed (*k* = 10) than in Fig. 3 (*k* = 1) (see Methods). Black curves with arrows show the system’s oscillatory trajectory. C. course of species biomass in the chemostat over a long duration for species in (B).

**Figure S4.**
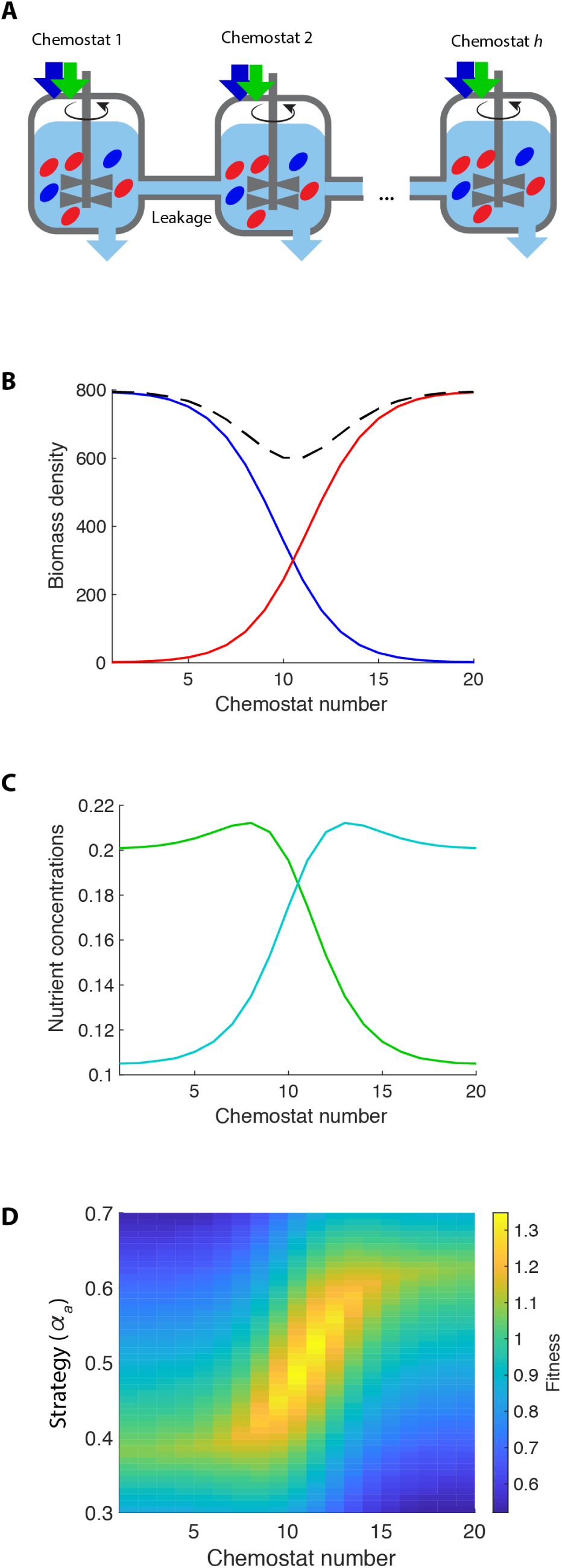
Steady-state spatial heterogeneity for linked chemostats. With initial seeding of two species, one at each of the two ends of a chain of chemostats, a steady-state gradient of species biomass density spontaneously emerges accompanied by a gradient of nutrient concentrations, even though the supply conditions and dilution rates are identical for all the chemostats. A. Schematic of *k*_tot_ linked chemostats exchanging medium and cells via leakage, described by Eqs. S34-S35. The two species in the chemostats (*Blue* and *Red*) are the same bistable pair as in Fig 4B and the leakage rate is *l* = 1. B. The species composition along 20 linked chemostats for the system in (A). Species colors correspond to those in Fig 4B, with species *Blue* having *α*_*a*_ = 0.35 and species *Red* having *α*_*a*_ = 0.65. The dashed black curve shows the sum of the two biomass densities. The initial condition was cell-free chemostats with a small amount of *Blue* added to Chemostat 1 and small amount of *Red* added to Chemostat 20. C. Concentrations along the 20 chemostats for nutrient *a* (green) and nutrient *b* (cyan) for system in (A). D. The fitness landscape along the chain of chemostats. The *x*-axis is the 20 linked chemostats, and the *y*-axis is the metabolic strategy represented by *α*_*a*_. Color indicates the growth rate of species adopting the given strategy in the *k*-th chemostat.

**Figure S5.**
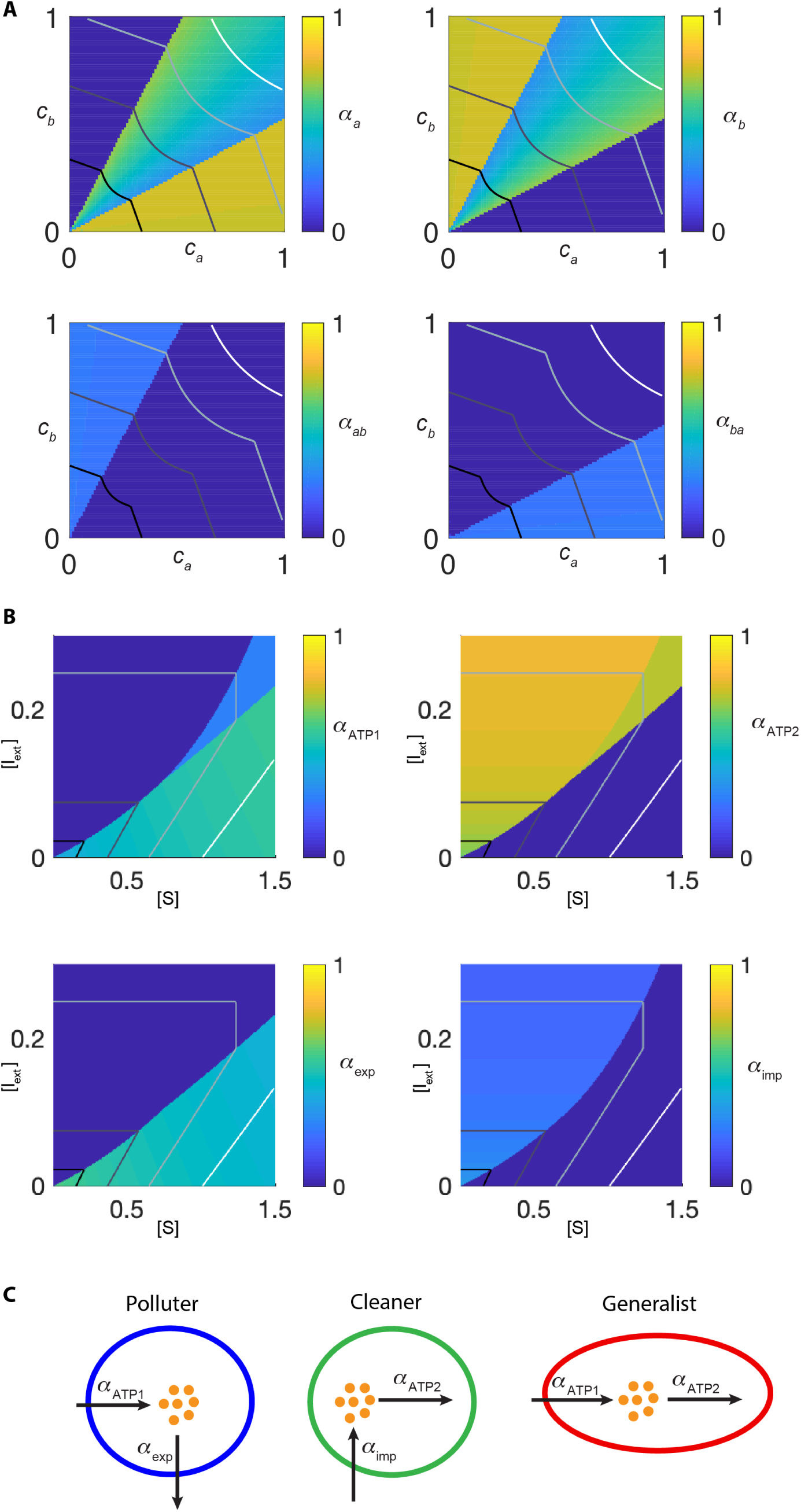
Maximizing strategies in chemical space. A. For each environment in the chemical space, the maximizing resource allocation strategies that maximize growth rates for the model in Fig 5A. Each strategy is represented by the four elements [*α*_*a*_, *α*_*b*_, *α*_*ab*_, *α*_*ba*_], and values for each element are shown by a heatmap. Black-to-white curves are the maximal growth contours for *d* = 0.1, 0.2, 0.3, 0.4. B. For each environment in the chemical space, the maximizing resource allocation strategies that maximize growth rates for the model in Fig 6A. Each strategy is represented by the four elements [*α*_ATP1_, *α*_ATP2_, *α*_exp_, *α*_imp_], and values for each element are shown by a heatmap. Black-to-white curves are the maximal growth contours for *d* = 0.2, 0.4, 0.6. C. Schematic representations of the three classes of maximizing strategies appearing in (B).

## REFERENCES

Armstrong, R. A., & McGehee, R. (1980). Competitive exclusion. The American Naturalist, 115(2), 151–170.

Bachmann, H., Bruggeman, F. J., Molenaar, D., dos Santos, F. B., & Teusink, B. (2016). Public goods and metabolic strategies. Current Opinion in Microbiology, 31, 109–115. doi: 10.1016/j.mib.2016.03.007

Bachmann, H., Molenaar, D., dos Santos, F. B., & Teusink, B. (2017). Experimental evolution and the adjustment of metabolic strategies in lactic acid bacteria. Fems Microbiology Reviews, 41, S201–S219. doi: 10.1093/femsre/fux024

Bajic, D., & Sanchez, A. (2019). The ecology and evolution of microbial metabolic strategies. Curr Opin Biotechnol, 62, 123–128. doi: 10.1016/j.copbio.2019.09.003

Bajic, D., Vila, J. C. C., Blount, Z. D., & Sanchez, A. (2018). On the deformability of an empirical fitness landscape by microbial evolution. Proc Natl Acad Sci U S A, 115(44), 11286–11291. doi: 10.1073/pnas.1808485115

Beardmore, R. E., Gudelj, I., Lipson, D. A., & Hurst, L. D. (2011). Metabolic trade-offs and the maintenance of the fittest and the flattest. Nature, 472(7343), 342–346. doi: 10.1038/nature09905

Blount, Z. D., Barrick, J. E., Davidson, C. J., & Lenski, R. E. (2012). Genomic analysis of a key innovation in an experimental Escherichia coli population. Nature, 489(7417), 513-+. doi: 10.1038/nature11514

Boer, V. M., Crutchfield, C. A., Bradley, P. H., Botstein, D., & Rabinowitz, J. D. (2010). Growth-limiting intracellular metabolites in yeast growing under diverse nutrient limitations. Mol Biol Cell, 21(1), 198–211. doi: 10.1091/mbc.E09-07-0597

Callahan, B. J., Fukami, T., & Fisher, D. S. (2014). Rapid evolution of adaptive niche construction in experimental microbial populations. Evolution, 68(11), 3307–3316. doi: 10.1111/evo.12512

Chase, J. M., & Leibold, M. A. (2003). Ecological niches: linking classical and contemporary approaches: University of Chicago Press.

D’Souza, G., Shitut, S., Preussger, D., Yousif, G., Waschina, S., & Kost, C. (2018). Ecology and evolution of metabolic cross-feeding interactions in bacteria. Nat Prod Rep, 35(5), 455–488. doi: 10.1039/c8np00009c

De Leenheer, P., Levin, S. A., Sontag, E. D., & Klausmeier, C. A. (2006). Global stability in a chemostat with multiple nutrients. Journal of Mathematical Biology, 52(4), 419–438. doi: 10.1007/s00285-005-0344-4

Doebeli, M. (2002). A model for the evolutionary dynamics of cross-feeding polymorphisms in microorganisms. Population Ecology, 44(2), 59–70. doi: DOI 10.1007/s101440200008

Escalante-Chong, R., Savir, Y., Carroll, S. M., Ingraham, J. B., Wang, J., Marx, C. J., & Springer, M. (2015). Galactose metabolic genes in yeast respond to a ratio of galactose and glucose. Proc Natl Acad Sci U S A, 112(5), 1636–1641. doi: 10.1073/pnas.1418058112

Friedman, J., Higgins, L. M., & Gore, J. (2017). Community structure follows simple assembly rules in microbial microcosms. Nature ecology & evolution, 1(5), 0109.

Goldford, J. E., Lu, N., Bajić, D., Estrela, S., Tikhonov, M., Sanchez-Gorostiaga, A., … Sanchez, A. (2018). Emergent simplicity in microbial community assembly. Science, 361(6401), 469–474.

Goyal, A., Dubinkina, V., & Maslov, S. (2018). Multiple stable states in microbial communities explained by the stable marriage problem. The ISME Journal. doi: 10.1038/s41396-018-0222-x

Goyal, S., Yuan, J., Chen, T., Rabinowitz, J. D., & Wingreen, N. S. (2010). Achieving optimal growth through product feedback inhibition in metabolism. PLoS computational biology, 6(6), e1000802.

Hardin, G. (1960). The competitive exclusion principle. Science, 131(3409), 1292–1297.

Huisman, J., van Oostveen, P., & Weissing, F. J. (1999). Species dynamics in phytoplankton blooms: incomplete mixing and competition for light. The American Naturalist, 154(1), 46–68.

Huisman, J., & Weissing, F. J. (1999). Biodiversity of plankton by species oscillations and chaos. Nature, 402(6760), 407–410. doi: 10.1038/46540

Huisman, J., & Weissing, F. J. (2001). Biological conditions for oscillations and chaos generated by multispecies competition. Ecology, 82(10), 2682–2695.

Ispolatov, I., Madhok, V., & Doebeli, M. (2016). Individual-based models for adaptive diversification in high-dimensional phenotype spaces. Journal of theoretical biology, 390, 97–105. doi: 10.1016/j.jtbi.2015.10.009

Kasting, J. F., & Siefert, J. L. (2002). Life and the evolution of Earth’s atmosphere. Science, 296(5570), 1066–1068. doi: 10.1126/science.1071184

Koffel, T., Daufresne, T., Massol, F., & Klausmeier, C. A. (2016). Geometrical envelopes: Extending graphical contemporary niche theory to communities and eco-evolutionary dynamics. Journal of theoretical biology, 407, 271–289.

Levin, S. A. (1970). Community equilibria and stability, and an extension of the competitive exclusion principle. The American Naturalist, 104(939), 413–423.

Li, S. H.-J., Li, Z., Park, J. O., King, C. G., Rabinowitz, J. D., Wingreen, N. S., & Gitai, Z. (2018). Escherichia coli translation strategies differ across carbon, nitrogen and phosphorus limitation conditions. Nature microbiology, 3(8), 939.

Liebermeister, W., Noor, E., Flamholz, A., Davidi, D., Bernhardt, J., & Milo, R. (2014). Visual account of protein investment in cellular functions. Proceedings of the National Academy of Sciences, 111(23), 8488–8493.

Long, C. P., & Antoniewicz, M. R. (2018). How adaptive evolution reshapes metabolism to improve fitness: recent advances and future outlook. Current Opinion in Chemical Engineering, 22, 209–215. doi: 10.1016/j.coche.2018.11.001

Luli, G. W., & Strohl, W. R. (1990). Comparison of growth, acetate production, and acetate inhibition of Escherichia coli strains in batch and fed-batch fermentations. Applied and environmental microbiology, 56(4), 1004–1011.

MacArthur, R. (1970). Species packing and competitive equilibrium for many species. Theoretical population biology, 1(1), 1–11.

Maharjan, R., Seeto, S., Notley-McRobb, L., & Ferenci, T. (2006). Clonal adaptive radiation in a constant environment. Science, 313(5786), 514–517. doi: 10.1126/science.1129865

Marsland III, R., Cui, W., & Mehta, P. (2019). The Minimum Environmental Perturbation Principle: A New Perspective on Niche Theory. arXiv preprint 1901.09673.

McGehee, R., & Armstrong, R. A. (1977). Some mathematical problems concerning the ecological principle of competitive exclusion. Journal of Differential Equations, 23(1), 30–52.

Metz, J. (2012). Adaptive dynamics.

Odum, E. P., & Barrett, G. W. (1971). Fundamentals of ecology (Vol. 3): Saunders Philadelphia.

Pfeiffer, T., & Bonhoeffer, S. (2004). Evolution of cross-feeding in microbial populations. The American Naturalist, 163(6), E126–E135.

Posfai, A., Taillefumier, T., & Wingreen, N. S. (2017). Metabolic Trade-Offs Promote Diversity in a Model Ecosystem. Physical Review Letters, 118(2). doi: 10.1103/PhysRevLett.118.028103

Roller, B. R. K., Stoddard, S. F., & Schmidt, T. M. (2016). Exploiting rRNA operon copy number to investigate bacterial reproductive strategies. Nature Microbiology, 1(11). doi:Unsp 16160 10.1038/Nmicrobiol.2016.160

Rosenzweig, R. F., Sharp, R., Treves, D. S., & Adams, J. (1994). Microbial evolution in a simple unstructured environment: genetic differentiation in Escherichia coli. Genetics, 137(4), 903–917.

Rueffler, C., Van Dooren, T. J. M., & Metz, J. A. J. (2004). Adaptive walks on changing landscapes: Levins’ approach extended. Theoretical Population Biology, 65(2), 165–178. doi: 10.1016/j.tpb.2003.10.001

Schertzer, E., Staver, A. C., & Levin, S. A. (2015). Implications of the spatial dynamics of fire spread for the bistability of savanna and forest. J Math Biol, 70(1-2), 329–341. doi: 10.1007/s00285-014-0757-z

Schirrmeister, B. E., de Vos, J. M., Antonelli, A., & Bagheri, H. C. (2013). Evolution of multicellularity coincided with increased diversification of cyanobacteria and the Great Oxidation Event. Proc Natl Acad Sci U S A, 110(5), 1791–1796. doi: 10.1073/pnas.1209927110

Scott, M., Gunderson, C. W., Mateescu, E. M., Zhang, Z. G., & Hwa, T. (2010). Interdependence of Cell Growth and Gene Expression: Origins and Consequences. Science, 330(6007), 1099–1102. doi: 10.1126/science.1192588

Smith, H. L., & Waltman, P. (1995). The theory of the chemostat: dynamics of microbial competition (Vol. 13): Cambridge university press.

Soliveres, S., Maestre, F. T., Ulrich, W., Manning, P., Boch, S., Bowker, M. A., … Allan, E. (2015). Intransitive competition is widespread in plant communities and maintains their species richness. Ecology letters, 18(8), 790–798. doi: 10.1111/ele.12456

Taillefumier, T., Posfai, A., Meir, Y., & Wingreen, N. S. (2017). Microbial consortia at steady supply. eLife, 6, e22644.

Tilman, D. (1980). Resources: a graphical-mechanistic approach to competition and predation. The American Naturalist, 116(3), 362–393.

Tilman, D. (1982). Resource competition and community structure: Princeton university press.

Van den Bergh, B., Swings, T., Fauvart, M., & Michiels, J. (2018). Experimental design, population dynamics, and diversity in microbial experimental evolution. Microbiol. Mol. Biol. Rev., 82(3), e00008–00018.

Wang, X., & Tang, C. (2017). Optimal growth of microbes on mixed carbon sources. arXiv preprint 1703.08791.

Wides, A., & Milo, R. (2018). Understanding the dynamics and optimizing the performance of chemostat selection experiments. arXiv preprint 1806.00272.

Zaman, S., Lippman, S. I., Zhao, X., & Broach, J. R. (2008). How Saccharomyces Responds to Nutrients. Annual Review of Genetics, 42, 27–81. doi: 10.1146/annurev.genet.41.110306.130206

Ziv, N., Brandt, N. J., & Gresham, D. (2013). The Use of Chemostats in Microbial Systems Biology. Jove-Journal of Visualized Experiments (80). doi:UNSP e50168 10.3791/50168

